# Molecular organization of autonomic, respiratory, and spinally-projecting neurons in the mouse ventrolateral medulla

**DOI:** 10.1101/2023.11.15.564801

**Authors:** Dana C. Schwalbe, Daniel S. Stornetta, Ruei-Jen Abraham-Fan, George M. P. R. Souza, Maira Jalil, Maisie E. Crook, John N. Campbell, Stephen B. G. Abbott

## Abstract

The ventrolateral medulla (VLM) is a crucial region in the brain for visceral and somatic control. It also serves as a significant source of synaptic input to the spinal cord. Experimental studies have shown that gene expression in individual VLM neurons is predictive of their function. However, the organizing principles of the VLM have remained uncertain. This study aimed to create a comprehensive dataset of VLM cells using single-cell RNA sequencing. The dataset was enriched with targeted sequencing of spinally-projecting and adrenergic/noradrenergic VLM neurons. Based on differentially expressed genes, the resulting dataset of 114,805 VLM cells identifies 23 subtypes of neurons, excluding those in the inferior olive, and 5 subtypes of astrocytes. Spinally-projecting neurons were found to be abundant in 7 subtypes of neurons, which were validated through in-situ hybridization. These subtypes included adrenergic/noradrenergic neurons, serotonergic neurons, and neurons expressing gene markers associated with pre-motor neurons in the ventromedial medulla. Further analysis of adrenergic/noradrenergic neurons and serotonergic neurons identified 9 and 6 subtypes, respectively, within each class of monoaminergic neurons. Marker genes that identify the neural network responsible for breathing were concentrated in 2 subtypes of neurons, delineated from each other by markers for excitatory and inhibitory neurons. These datasets are available for public download and for analysis with a user-friendly interface. Collectively, this study provides a fine-scale molecular identification of cells in the VLM, forming the foundation for a better understanding of the VLM’s role in vital functions and motor control.

## Introduction

The ventrolateral medulla (VLM) is a brain region that is critical for the regulation of autonomic function, respiratory control, sleep-wake behavior, cranial motor function and locomotion (Arber and Costa, 2022; Coverdell et al., 2022; Del Negro et al., 2018; Guyenet, 2014; Kleinfeld et al., 2023; Scammell et al., 2017). In adult mammals, the VLM is characterized by interlacing networks of fiber bundles and lacks easily demarcated cyto-architectural boundaries (Watson et al., 2019). However, neurophysiological and neuroanatomical studies have identified functionally-specialized groups of neurons in the VLM associated with unique molecular profiles (Coverdell et al., 2022; Guyenet et al., 2013; Veerakumar et al., 2022; Yackle, 2023) and functionally specialized astrocytes (Kasymov et al., 2013). As these studies have shown, establishing the transcriptional landscape of the VLM is important for understanding the VLM’s role in vital functions. However, our understanding of the molecular landscape of the VLM is still limited, posing a challenge for experimental studies. Single-nuclei RNA sequencing (snRNA-seq) parses cellular diversity based on the genome-wide expression profiles of individual cells and groups them by similarity into candidate cell types. Applying this approach to discrete brain regions in mice has revealed remarkable diversity of cell types, raising hypotheses about their functional roles, and providing genetic access points for their further investigation (e.g., (Bakken et al., 2018; Habib et al., 2016; Lake et al., 2016; Liu et al., 2023)). A comprehensive molecular database of VLM cell types does not exist, although several studies have applied single-cell or single nuclei RNA-seq to particular cell populations within the VLM (Bachmutsky et al., 2020; Beine et al., 2022; Coverdell et al., 2022; Hayes et al., 2017; Veerakumar et al., 2022). We therefore used high-throughput snRNA-seq to systematically characterize the molecular cell types of the mouse VLM. Furthermore, we used anatomically- and genetically-targeted cell-sorting of spinally-projecting and adrenergic/noradrenergic VLM neurons to identify and characterize neurons that provide input to neurons in the spinal cord involved in sympathetic and locomotor function.

## Results

### Transcriptional Analysis Confirms Major Cell Classes and Uncovers Astrocyte Heterogeneity

To classify cell types of the VLM by their gene expression profiles, we dissected VLM tissue from adult male and female mice for single-nuclei RNA-sequencing (snRNA-seq). We used naive C57Bl6J mice and H2B-TRAP(mCherry) Cre-reporter mice which were either crossed with DBH-Cre mice, resulting in H2B-mCherry expression in VLM adrenergic/noradrenergic neurons (DBH^TRAP^ mice; Figure 1A-C), or injected in the thoracic spinal cord with a retrograde AAV-Cre, resulting in H2B-mCherry expression in spinally-projecting neurons (Spinal^TRAP^ mice) (Figure 1D-G). In DBH^TRAP^ mice, nuclear H2B-mCherry expression was observed in 76 ± 8% of adrenergic/noradrenergic VLM neurons identified by expression of *Dbh* by RNA fluorescent *in situ* hybridization (FISH) (Figure 1B). Furthermore, nuclear H2B-mCherry expression was observed in 84 ± 8% tyrosine hydroxylase (TH) expressing VLM neurons, determined by immunofluorescence (Figure 1C). Importantly, *Dbh* expression was detected in all cells that expressed H2B-mCherry (based on 313 H2B-mCherry+ cells in 3 cases) and 89 ± 3% of H2B-mCherry+ cells expressed TH. Hence, DBH^TRAP^ mice provide a selective and efficient targeting approach to capture adrenergic/noradrenergic VLM neurons. In Spinal^TRAP^ mice, we observed H2B-mCherry expression in neurons in the VLM and ventromedial medulla (VMM) (Figure 1D-G), including a subpopulation of TH+ neurons in the rostral VLM (Figure 1G). To control for the possibility that cells were transduced due to the leakage of AAV into the CSF, we pipetted 1 µL of AAV onto the exposed surface of the thoracic spinal cord followed by wound irrigation and closure. This control procedure did not produce any detectable H2B-mCherry expression in the brainstem.

**Figure 1:**
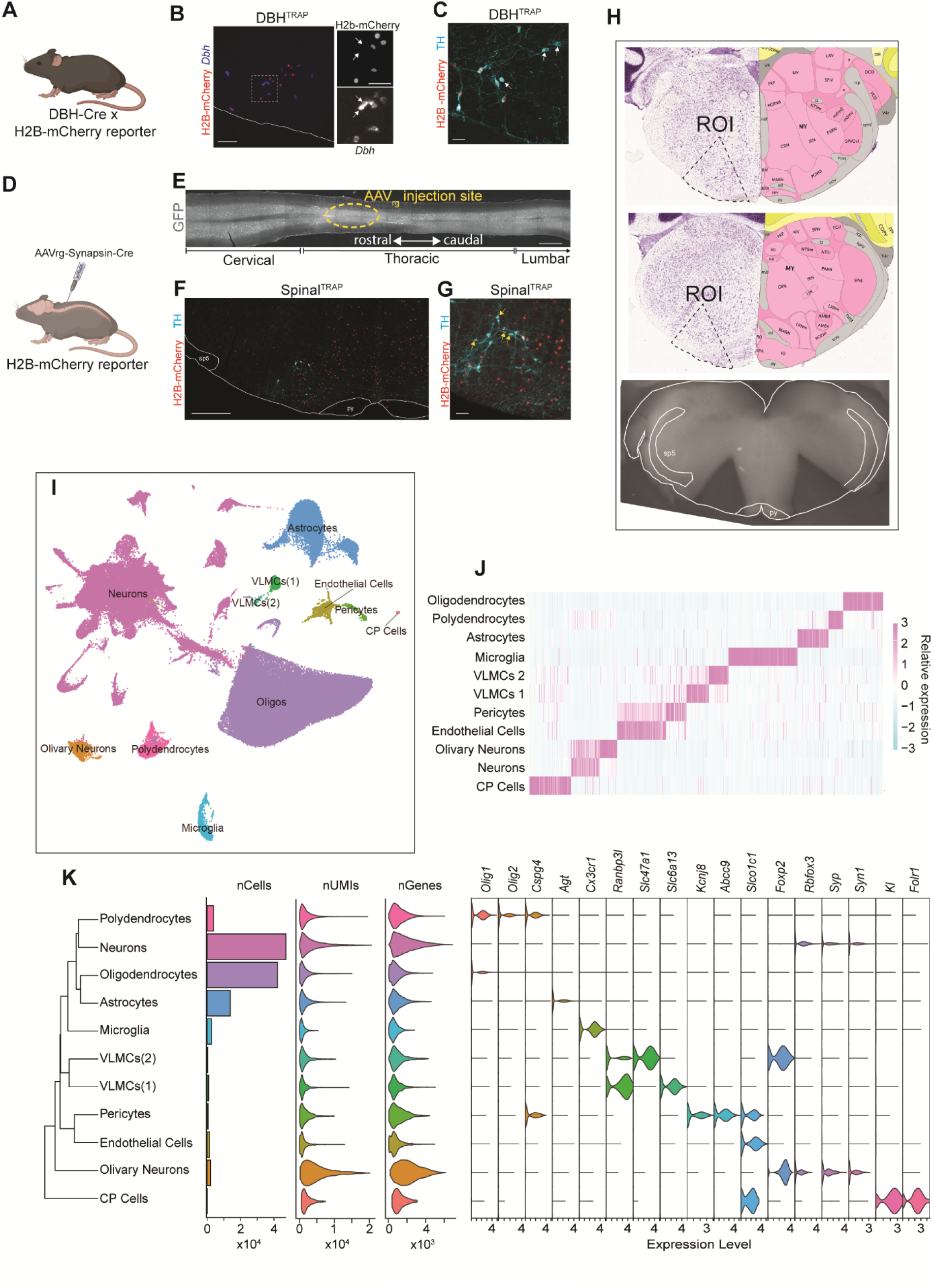
Approach for single-nuclei RNA-seq of the ventrolateral medulla (VLM) A. Genetic approach to label *Dbh*-expressing neurons for collection, *i.e.*, generation of DBH^TRAP^ mice. Figure made with BioRender. B. Expression of H2B-mCherry in the VLM of DBH^Trap^ mice colocalizes with *Dbh*+ neurons by RNA fluorescence *in situ* hybridization (FISH). White arrows identify *Dbh*+ neurons that did not contain a H2B-mCherry positive nuclei. Scale bars: 100 µm left panel and 50 µm for right panels C. Expression of H2B-mCherry in the VLM of DBH^TRAP^ mice colocalizes with TH immunofluorescence. White arrows identify TH+ neurons that did not contain a H2B-mCherry positive nuclei. Scale bar-50 µm D. Approach to label spinally-projecting neurons for collection, *i.e.*, generation of Spinal^TRAP^ mice. Figure made with BioRender. E. Coronal section of the spinal cord from a Spinal^TRAP^ mouse showing the location and approximate spread of AAV injections. F. Expression of H2B-mCherry in the VLM of Spinal^TRAP^ mice is distributed in the ventrolateral and ventromedial medulla. Scale bar-500 µm G. Spinal^TRAP^ cells in the rostral VLM express TH. Yellow arrows identify TH+ neurons that have H2B-mCherry positive nuclei. Scale bar-50 µm H. Region of the medulla collected for analysis. Upper and middle panels show the region of interest superimposed on Nissl (left) and anatomical annotations (right) from the Allen Mouse Brain Atlas and Allen Reference Atlas – Mouse Brain. Allen Mouse Brain Atlas, mouse.brain-map.org and atlas.brain-map.org. Lower panel shows the remaining tissue after two bilateral wedges of tissue were collected from a region encompassing the VLM for tissue processing. I. Uniform Manifold Approximation and Projection (UMAP) of all cells, labeled according to cell type. J. Statistically significant differentially expressed genes across all cell types. K. Dendrogram, number of cells, unique molecular identifiers, genes detected, and the expression of cell type marker genes for cell types shown in I.

To prepare samples for snRNA-seq of the VLM, we collected 1 mm thick transverse sections of the medulla between -6 mm to -8 mm caudal from bregma. We then dissected bilateral ‘wedges’ of the VLM bounded medially by the pyramids and laterally by the spinal trigeminal tract with the tip of the wedge in the intermediate reticular nucleus (Figure 1H). Care was taken to avoid collecting midline raphe nuclei (raphe pallidus and obscurus) and the trigeminal region. Based on reference anatomical atlases, the cells collected for sequencing likely originated from regions including: paragigantocellular reticular nucleus; parapyramidal nucleus; nucleus ambiguus; facial nucleus; lateral reticular nucleus; magnocellular reticular nucleus; gigantocellular reticular nucleus; intermediate reticular nucleus; and the inferior olive (Figure 1H).

We isolated cell nuclei from the dissected VLM tissue and profiled their genome-wide mRNA content by snRNA-seq. Nuclei from DBH^TRAP^ and Spinal^TRAP^ tissue samples were sorted by flow cytometry to separately purify single mCherry-positive and -negative cell nuclei, whose poly-adenylated RNA were then processed in parallel into sequencing libraries. After sequencing the libraries and quantifying genome-wide expression values, we filtered out low-quality transcriptomes and putative cell doublets (see Methods and workflow illustrated in Supplementary Figure 1). This procedure left 114,805 single-nuclei transcriptomes (∼cells) for further analysis. Annotated Seurat code for the dataset can be accessed through Zenodo (record number: 10048289).

We integrated the cells across six sample batches (Supplemental figure 2A-C), clustered them based on similarity of expression of high-variance genes, and annotated each cluster according to its expression of canonical cell class marker genes (Figure 1I-K). Clusters of neurons, identified by their expression of genes encoding NeuN (*Rbfox3*), Synaptophysin (*Syp*), and Synapsin 1 (*Syn1*), comprised the largest proportion of cells at 41.0%. A smaller cluster of neurons (*Syn1*, *Rbfox3*) distinguished by high expression *Foxp2* likely originated from the inferior olive (*Foxp2*+ neurons; 1.8% of cells; Supplemental figure 2C,D). Oligodendrocytes, marked by their expression of the oligodendrocyte transcription factor gene *Olig1*, made up the second largest cluster at 36.6% of all cells. The relative abundance of oligodendrocytes was expected given the high white matter content of the VLM. Astrocytes, identified by their expression of angiotensinogen (*Agt*), made up the third largest cell cluster (12.0% of cells). Interestingly, reclustering the astrocytes apart from the other cell types revealed five molecularly-distinct subtypes (Supplemental Figure 2E). Polydendrocytes, also known as oligodendrocyte precursor cells (OPCs), were distinguished by their expression of the oligodendrocyte transcription factor genes *Olig1* and *Olig2* and the chondroitin sulfate proteoglycan 4 (NG2) gene *Cspg4* and represented 3.3% of all cells. Approximately 2.3% of the cells formed a macrophage cluster, containing mostly microglia (*P2ry12*+) and a relatively smaller number of perivascular macrophages (*Mrc1*+). The remainder of the dataset comprised two distinct clusters of vascular and leptomeningeal cells (*Ranbp3l*+; 1.0% of cells) and one cluster each of pericytes (*Kcnj8*+, *Abcc9*+; 0.5% of cells), endothelial cells (*Slco1c1*+; 1.3% of cells), and choroid plexus cells (CP; *Kl*+, *Folr1*+; 0.1% of cells). Overall, our molecular taxonomy of VLM cells includes all expected cell classes and reveals an unexpected degree of molecular heterogeneity among VLM astrocytes.

### Subclustering Neurons Reveals 23 Candidate Molecular Subtypes

To better resolve the diversity of VLM neurons, we subsetted all neurons except those putatively from the inferior olive and reclustered them apart from the non-neuronal cells. After removing potential doublets and molecularly ambiguous cells (see Methods and Supplementary Figure 1), we grouped 15,342 neurons into 23 clusters (Figure 2A). On average (+/- standard deviation), we detected 2,277 +/- 1,213 genes in each neuron, based on 4,806 +/- 3,760 transcripts per neuron (Figure 2B).

**Figure 2:**
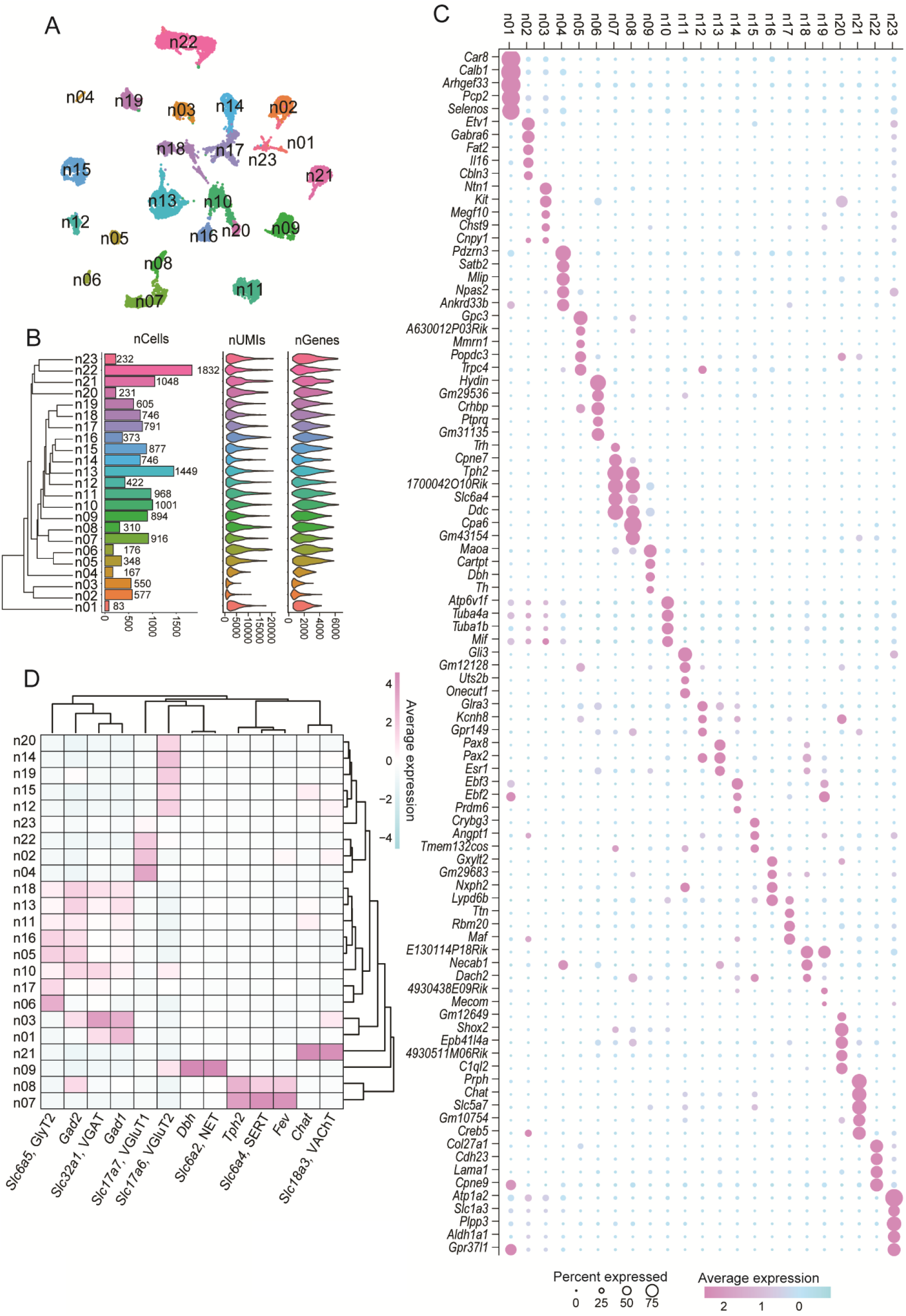
Diversity of neurons in the VLM. A. UMAP of all neurons, labeled by cluster ID B. Dendrogram showing neuron cluster relatedness; numbers of cells per cluster, UMI per cell, and genes detected per cell for each of the neuron clusters shown in A. C. Dot plot of top differentially expressed genes for neuron clusters shown in A D. Heat map of neurotransmitter phenotype related genes for neuron clusters shown in A.

Comparing gene expression across neuronal clusters revealed candidate marker genes for each cluster (Figure 2C). To identify the anatomical origins of each neuronal cluster, we cross-referenced its top marker genes with publicly available ISH data from the Allen Brain Atlas (Lein et al., 2007). Table 1 provides a summary of the putative anatomical origin of each population based on this analysis. To predict the neurochemical phenotype of each neuron cluster, we assessed the expression of genes involved in neurotransmitter synthesis and cell identity (Figure 2D): excitatory neurons (*Slc17a7*/VGluT2, *Slc17a6*/vGluT2); inhibitory neurons (*Slc32a1*/VGAT, *Slc6a5*/GlyT2, *Gad1*/GAD67, *Gad2*/GAD65); cholinergic neurons (*Chat*/ChAT, *Slc18a3*/VAChT); serotonergic neurons (*Slc6a4*/SERT1, *Tph2*/Tryptophan Hydroxylase 2, *Fev*/Pet-1); and adrenergic/noradrenergic neurons (*Dbh*/Dopamine Beta-Hydroxylase, *Slc17a6*/vGluT2 and *Slc6a2*/NET). This analysis identifies 9 clusters of inhibitory neurons; 7 clusters of excitatory neurons; 2 clusters of serotonergic neurons (n07, n08); 1 cluster of adrenergic/noradrenergic neurons (n09) and 1 cluster of cholinergic neurons (n21). Two clusters (n10, n23) expressed markers for both excitatory and inhibitory neurons but rarely in the same cells, suggesting these clusters are heterogeneous.

**Table 1.** Presumptive location of neuron cluster based on differentially expressed genes and neurotransmitter phenotype.

| Neuron cluster id | Differentially expressed genes with traceable expression in ABA | Enriched transcription factors | Neurotransmitter phenotype | Putative location |
| --- | --- | --- | --- | --- |
| 1 | *Pcp2, Arhgap31, Foxp2* | *Lhx1, Foxp2* | inhibitory | unclear |
| 2 | *Chn2, Etv1, il16* | *Neurod1* | excitatory (VGLUT1) | unclear |
| 3 | *Kit, Pvalb* |  | inhibitory | MARN |
| 4 | *Gabra4, slc17a7* |  | excitatory (VGLUT1) | unclear |
| 5 | *Gpc3, Crhbp, Trpc4* | *Gata3* | inhibitory | PGRN |
| 6 | *Crhbp* | *Gata3* | inhibitory | PGRN/MARN |
| 7 | *Tph2, Trh, Cpne7* | *Lmx1b* | serotonergic | PPY/MARN |
| 8 | *Tph2, Cpa6* | *Lmx1b* | serotonergic | PPY/MARN |
| 9 | *Cartpt, Dbh* | *Tfap2b* | noradrenergic/glutamatergic (VGLUT2) | PGRN |
| 10 | *none* | - | unclear | unclear |
| 11 | *Gli3, Uts2b* |  | inhibitory | PGRN |
| 12 | *Cdh9, Glra3, Trpc4* | *Pax2* | glutamatergic (VGLUT2) | PGRN |
| 13 | *Pax8, Pax2* | *Dmbx1, Pax2* | inhibitory | PGRN |
| 14 | *none* | - | glutamatergic (VGLUT2) | unclear |
| 15 | *Crybg3* |  | glutamatergic (VGLUT2) | PGRN |
| 16 | *Gxylt2, Lypd6b* | *Gata3* | inhibitory | PGRN |
| 17 | *none* | - | inhibitory | unclear |
| 18 | *none* | *Pax2* | inhibitory | unclear |
| 19 | *none* | - | glutamatergic (VGLUT2) | unclear |
| 20 | *C1ql2, Shox2* | *Lhx4, Lhx, Shox2* | glutamatergic (VGLUT2) | MARN |
| 21 | *Prph, Chat* | *Isl1* | Cholinergic | AMB/VII |
| 22 | *Cdh23, Col12a1* | - | glutamatergic (VGLUT1) | LRT |
| 23 | *none* | - | unclear | unclear |

### Spinal^TRAP^ Classifies Seven Molecular Subtypes of Bulbospinal Neurons

To identify neurons in our snRNA-seq dataset that innervate the spinal cord, we tracked Spinal^TRAP^ cells throughout the analysis and found that these cells distributed across several neuron clusters but were especially enriched in n06-n10, n16, and n20 (Figure 3A-C). Amongst these were serotonergic neurons (n07 and n08) and adrenergic/noradrenergic neurons (n09), both of which are a source of innervation for sympathetic preganglionic neurons in the spinal cord (Bowker et al., 1981; Tucker et al., 1987). Clusters n06, n16, n20 expressed novel marker transcripts that were detectable by ISH in the VLM and VMM (Figure 3D). The excitatory cluster n20 was marked by *C1ql2,* a gene expressed at high levels in VMM neurons, more specifically the ventral part of the gigantocellular nucleus and medial portion of the lateral paragigantocellular nucleus. The location and morphology of *C1ql2+* neurons (Figure 3D left and middle panels), neurotransmitter phenotype (Figure 2D) and transcription factors (Figure 3E) of cells in this cluster suggest that n20 represents excitatory reticulospinal neurons involved in locomotion and limb control (Bouvier et al., 2015; Bretzner and Brownstone, 2013; Capelli et al., 2017; Cepeda-Nieto et al., 2005; Esposito et al., 2014; Leiras et al., 2022). The other two populations of bulbospinal neurons (n06 and n16) in our dataset were inhibitory. Cluster n16 may represent a population of bulbospinal inhibitory neurons that are broadly distributed in the VLM based on the expression of marker genes (*Lypd6b*) (Figure 2C and Figure 3D, right panel). This cluster also expressed GlyT2 (*Scl6a5*) and VGAT (*Slc32a1*) (Figure 2D), similar to neurons identified in rats with a putative sympathetic function (Schreihofer et al., 1999; Stornetta et al., 2004). The other inhibitory bulbospinal cluster was defined by high expression of *Crhbp* (Figure 2C), which encodes a protein involved in the negative regulation of corticotropin releasing hormone, CRH (Potter et al., 1992, 1991). We mapped the distribution of bulbospinal *Crhbp* neurons by combining axonal tracing from the spinal cord with FISH. Neurons expressing *Crhbp* were located in both the VLM and VMM, with *Crhbp*+ cells in the VMM tending to have larger cell bodies than those in the VLM (Figure 3F). Bulbospinal *Crhbp+* neurons were located primarily in the ventral part of the gigantocellular nucleus and medial lateral paragigantocellular nucleus (Figure 3F,G), suggesting that that expression of *Crhbp* in the VMM may identify inhibitory neurons involved in behavioral arrest (Capelli et al., 2017).

**Figure 3.**
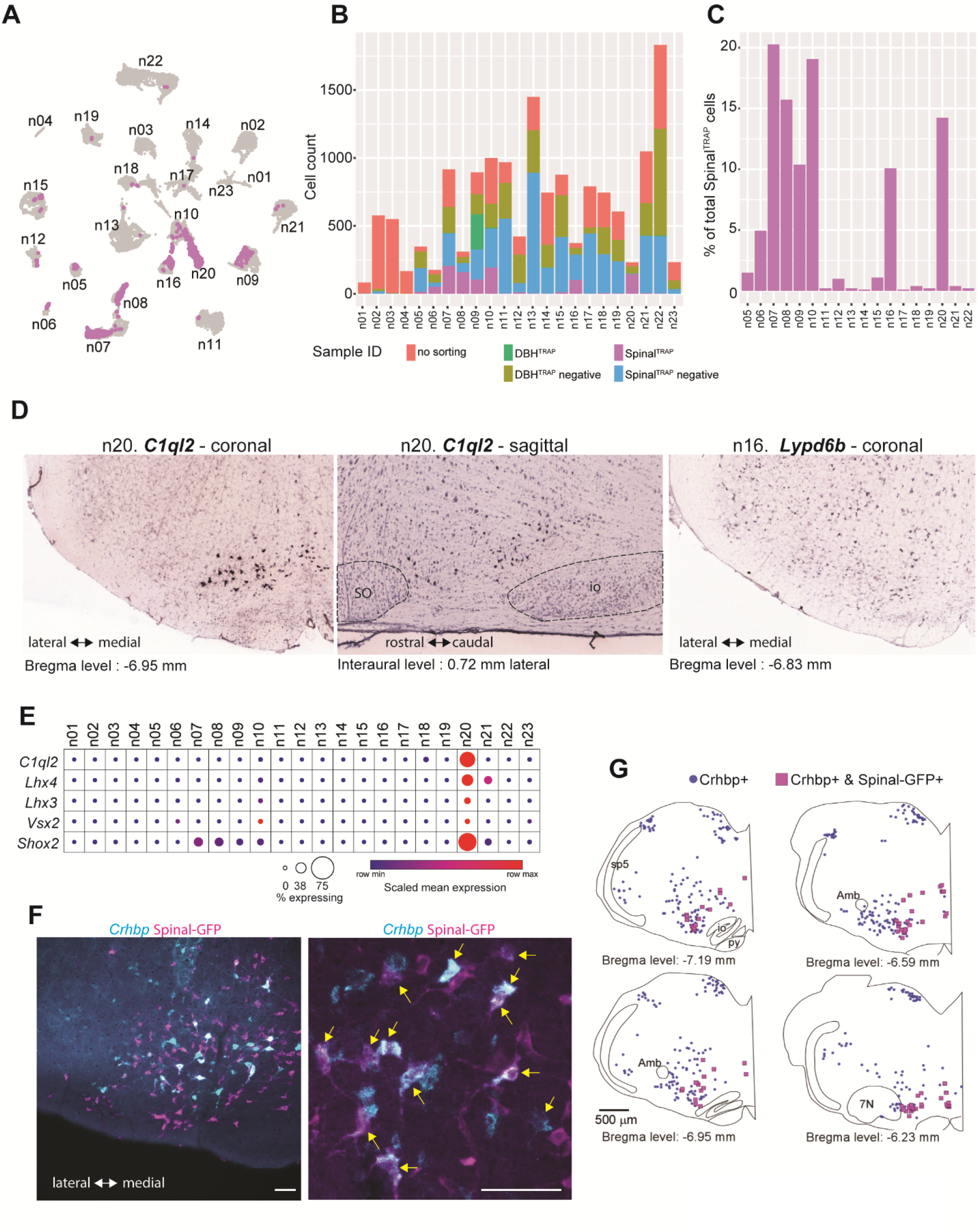
Spinal^TRAP^ identifies clusters of spinally projecting neurons. A. All-neuron UMAP (from Figure 2A) colored to show the distribution of Spinal^TRAP^ neurons B. Sample ID composition of neuron clusters. Note that clusters containing Spinal^TRAP^ neurons were integrated in clusters with cells from other samples. C. Distribution of Spinal^TRAP^ samples across neuron clusters, shown as a proportion of the total number of Spinal^TRAP^ cells in the dataset. D. Expression of *C1ql2* (https://mouse.brain-map.org/gene/show/86401) and *Lypd6b* (http://mouse.brain-map.org/experiment/show/77332684) in the VLM. E. Dot Plot of differentially expressed genes in n20 (*C1ql2* and *Shox2*) and genes expressed by neurons in the ventromedial medulla that are involved in motor control (*Lhx4*, *Lhx3*, and *Vsx2*/*Chx10;* see text for references). F. Expression of *Crhbp* by FISH in the VLM and overlap with spinally-projecting neurons (Spinal-GFP). Yellow arrows in the right panel indicate expression of *Crhbp* in spinally-projecting neurons. The images in the left and right panel are from different tissues. Scale bars-100 µm G. Distribution of *Crhbp* neurons in the medulla identifying the location where *Crhbp* was found in spinally-projecting neurons.

### Molecular Identification of VLM Respiratory Neurons

The VLM contains neuronal circuits that underlie respiratory and orofacial motor activity patterns (Kleinfeld et al., 2023; Smith, 2022). We reasoned that these networks might be represented in our dataset as one or more discrete neuron clusters. We examined the distribution of genes known to be expressed by respiratory-related neurons (*Tacr1*, *Sst*, *Oprm1*, *Penk)* (Bachmutsky et al., 2020; Cui et al., 2016; Gray et al., 1999; Stornetta et al., 2003b; Stornetta, 2008), which were enriched in several clusters in our snRNA-seq dataset but overlapped consistently in clusters n05 and n12 (Figure 4A). Interestingly, *Sst*, which is expressed in both inhibitory and excitatory respiratory-related neurons (Cui et al., 2016; Sherman et al., 2015; Stornetta et al., 2003a; Tan et al., 2008; Tupal et al., 2014) was enriched in n05, a cluster of inhibitory neurons, and expressed at relatively high levels in a small subset (∼5%) of neurons in n12, a cluster of excitatory neurons. Furthermore, *Cdh9*, a putative marker of excitatory respiratory-related neurons in mice (Vagnozzi et al., 2022; Yackle et al., 2017) was enriched in excitatory clusters n12, n16 and n18 (Figure 4A). Reelin, which is expressed in respiratory-related neurons (Tan et al., 2012), was also enriched in n05, n12, along with 7 other clusters. *Pax2*, which is expressed by many excitatory, but not inhibitory, respiratory-related neurons (Wu et al., 2017; Xia et al., 2022) was enriched in n12 and n18 supporting the possibility that these clusters contain excitatory respiratory-related neurons. Finally, neurons regulating diaphragm activity are reported to be enriched with *Cdh6* and *Cdh10,* in addition to *Cdh9,* in embryonic mouse medulla (Vagnozzi et al., 2022). In our snRNA-seq dataset based on adult mice, *Cdh6* and *Cdh10* expression was enriched in multiple clusters, suggesting that these genes do not effectively discriminate between respiratory neurons and other neurons in the VLM in adult mice. Altogether, the distribution of genes associated with respiratory-related neurons in our snRNA-seq dataset indicates clusters n05 and n12 are most likely to contain neurons involved in generating respiratory motor activity. The expression of marker genes for clusters n05 and n12 (*Gpc3* and *Glra3*, respectively) were localized to the VLM and VMM (Figure 4B,C). *Crhbp,* which was enriched in n05, is expressed by primarily non-bulbospinal neurons in respiratory-related regions of the VLM (Figure 3F). Also, both n05 and n12 had relatively high expression of *Trpc4*, which was also evident in cells localized to the respiratory column (Figure 4D). Of note, a relatively small number of Spinal^TRAP^ neurons was present in both n05 and n12, whereas n16 had a relatively large number of Spinal^TRAP^ neurons (Figure 3B,C). Interestingly, n16 also expressed markers predicted to be enriched phrenic premotor neurons in the rostral ventral respiratory group (*Penk* and *Pax2*) (Stornetta et al., 2003b; Wu et al., 2017). However, the primary neurotransmitter of cells in n16 is likely to be inhibitory based on the complement of vesicular transporters expressed in this cluster (Figure 4A), contrary to expectations for phrenic premotor neurons (Stornetta et al., 2003b; Wu et al., 2017). Finally, we examined our dataset for genes expressed by central chemoreceptor neurons in the retrotrapezoid nucleus (RTN), which regulate breathing in relation to brain pH (Guyenet et al., 2019). In mice, there are 600-700 RTN neurons, which can be distinguished from neighboring neurons by the expression of *Nmb* and *Gal* (Shi et al., 2017; Souza et al., 2023). In our dataset, *Nmb* and *Gal* were detected in only a few cells, and only co-detected in 4 cells across the entire dataset. This shows that RTN neurons do not form a unique cluster in our dataset, which may be attributed to these cells being a relatively small population of cells in the VLM.

**Figure 4.**
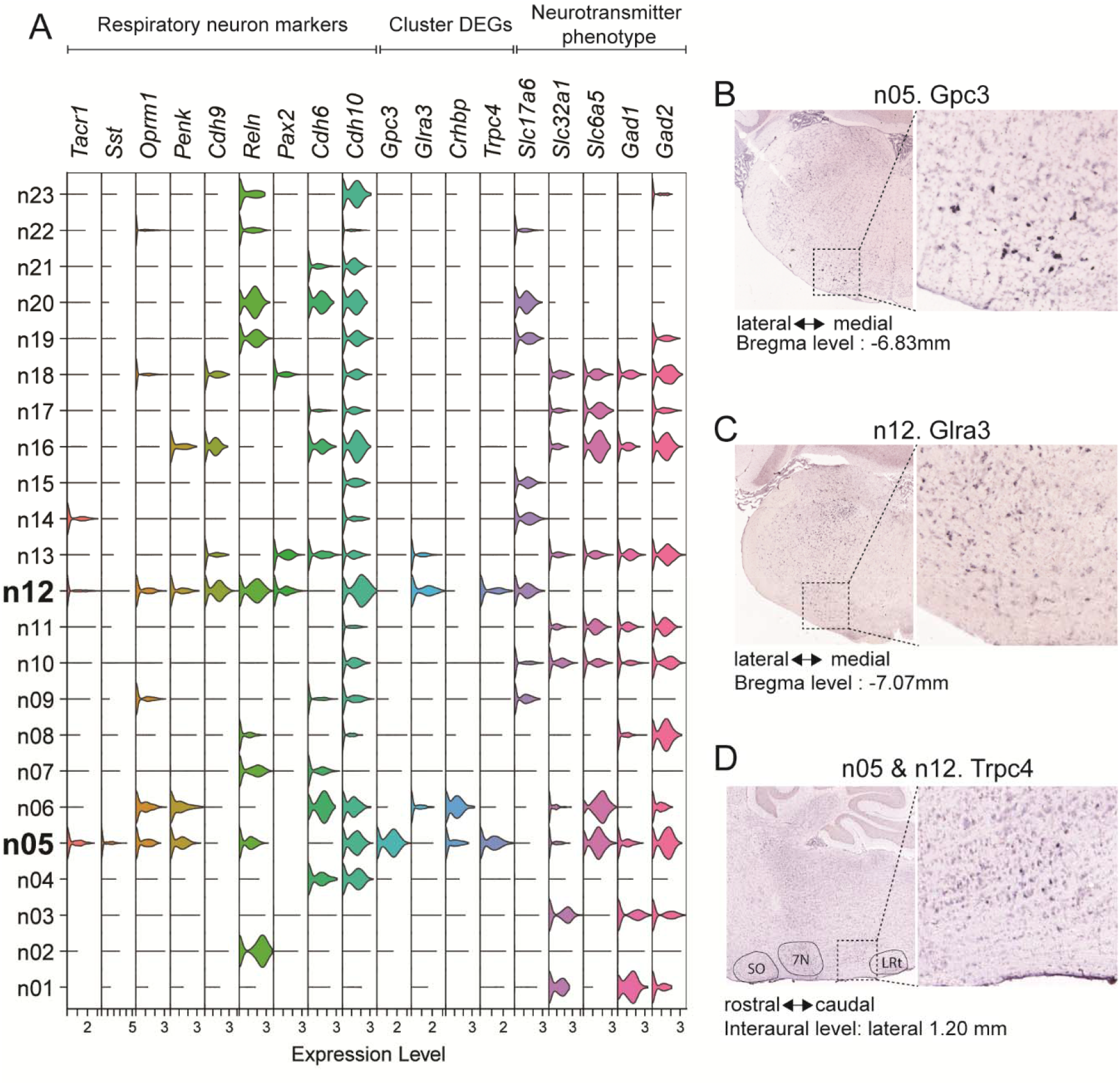
Identification of novel marker genes for VLM respiratory neurons. A. Violin plot of marker genes enriched for respiratory neurons (see text for references) and for neurotransmitter phenotypes, and differentially expressed genes for neuron cluster n05 and n12. B. Expression of *Gpc3* (http://mouse.brain-map.org/experiment/show/71020431) in the VLM from Allen Mouse Brain Atlas. C. *Glra3* (https://mouse.brain-map.org/experiment/show/73788474) in the VLM from Allen Mouse Brain Atlas. D. *Trpc4* (http://mouse.brain-map.org/experiment/show/1306) in the VLM from Allen Mouse Brain Atlas.

### Molecular Subtyping of Serotonergic Neurons

Single-cell transcriptomic analysis has revealed molecular diversity among serotonin neurons in the distinct raphe serotonergic cell groups (Huang et al., 2019; Okaty et al., 2015; Ren et al., 2019; Spaethling et al., 2014). To learn whether similar heterogeneity exists among the parapayramidal 5HT neurons, we sub-clustered the 5HT neurons in our snRNA-seq dataset. First, we selected neuron clusters expressing the phenotypic markers of 5HT cells (Figure 5A). These genes were restricted to neuron clusters n07 and n08, which differed from each other in their expression of *Trh*, *Cpne7*, *Cpa6*, and *Gm43154* (Figure 5A). Transcripts for marker genes of n07 and n08 were detected in both the parapyramidal region and midline raphe (pallidus and obscurus) (Figure 5B,C) and both n07 and n08 had a similar proportion of Spinal^TRAP^ neurons. This suggests that n07 and n08 represent molecularly distinct but anatomically intermixed subpopulations of 5HT neurons. Sub-clustering of 1226 cells from n07 and n08 resulted in 6 distinct sub-groups which differed in their expression of many novel marker genes (Figure 5D,F), as well as genes that differentiate 5HT neurons elsewhere in the brain (Figure 5G) (Huang et al., 2019; Okaty et al., 2015; Schneeberger et al., 2022; Xiao et al., 2021). Of note, Spinal^TRAP^ neurons were common in 5HT sub-clusters n07/08.s02, n07/08.s03 and n07/08.s04 (Figure 5E, Table 2). Collectively, this analysis indicates that the parapyramidal 5HT cell group comprises distinct populations of spinally-projecting neurons defined by a combination of newly identified marker genes as well as genes that discriminate 5HT neurons in other brain regions.

**Figure 5:**
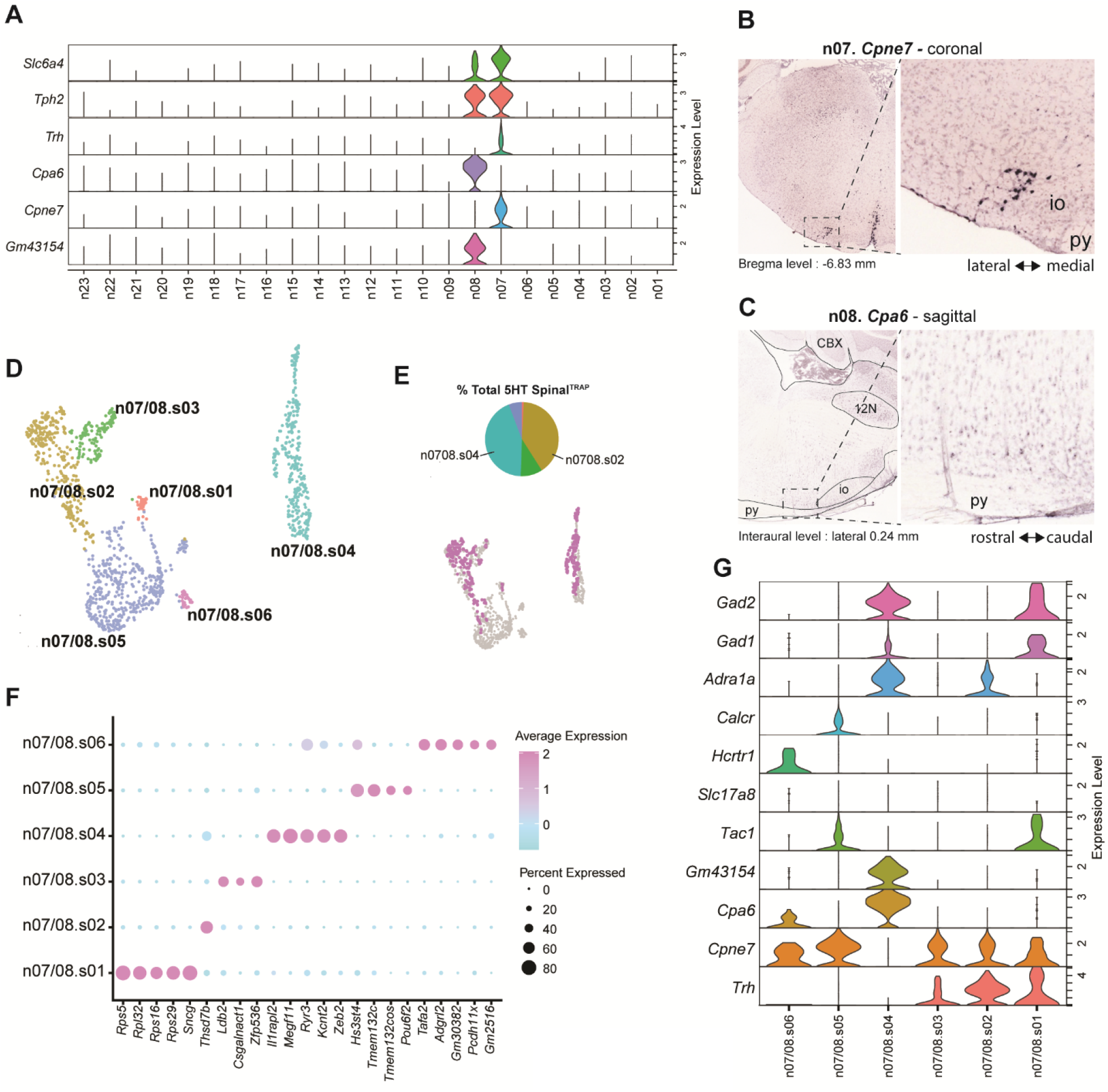
Identification of subtypes of serotonergic neurons in the parapyramidal region. A. Violin plot of phenotypic genes for serotonergic neurons (Slc6a4, Tph2) and expression of cluster marker genes for 5HT clusters n07 and n08 in ‘all-neuron’ clusters. B. Expression of Cpne7 in VLM from Allen Mouse Brain Atlas (http://mouse.brain-map.org/experiment/show/73817432). C. Expression of Cpa6 in VLM from Allen Mouse Brain Atlas (http://mouse.brain-map.org/experiment/show/75197606). D. UMAP generated by subclustering n07 and n08 based on high-variance genes E. Pie chart showing the proportion of 5HT SpinalTRAP samples in each sub-cluster and UMAP from panel D is colored to show the distribution of SpinalTRAP samples. Values for pie chart provided in table 2. F. Differentially expressed genes for 5HT neuron subclusters. G. Expression of genes that are differentially expressed in 5HT neurons in other brain regions.

**Table 2.** Values for pie charts in. **figure 5**

|  | Total |  | Spinal<sup>TRAP</sup> cells |  |
| --- | --- | --- | --- | --- |
| 5HT subclusters | no. of cells | % Total | no. of cells | % Total |
| n0708.s01 | 36 | 2.9 | 3 | 0.8 |
| n0708.s02 | 297 | 24.2 | 146 | 40.1 |
| n0708.s03 | 124 | 10.1 | 35 | 9.6 |
| n0708.s04 | 308 | 25.1 | 159 | 43.7 |
| n0708.s05 | 427 | 34.8 | 21 | 5.8 |
| n0708.s06 | 34 | 2.8 | 0 | 0.0 |
| Total | 1226 |  | 364 |  |

### Molecular Subtyping of Noradrenergic and Adrenergic Neurons

Adrenergic/noradrenergic neurons in the ventrolateral medulla regulate many important aspects of physiology, including the sympathetic control of blood pressure, glucose homeostasis, immune function, and the neuroendocrine responses to inflammation and stress (Guyenet et al., 2013). Our snRNA-seq analysis identified cluster n09 as adrenergic/noradrenergic neurons based on expression of *Cartpt*, *Dbh*, *Th*, and *Maoa* (Figure 6A). However, we noted that other markers for catecholaminergic cells were heterogeneously expressed among the neurons in cluster n09. For example, *Pnmt*, a marker for adrenergic neurons, and *Slc6a2*, the norepinephrine transporter, were expressed mostly by different cells in the n09 feature plot (Figure 6B). This observation aligns with the established expression patterns of *Pnmt* and *Slc6a2* expression in adrenergic/noradrenergic VLM neurons in rats (Guyenet et al., 2013). Similarly, we also noted that Spinal^TRAP^ neurons were not evenly distributed within cluster n09 (Figure 6B). Therefore, we sub-clustered n09 to further characterize the heterogeneity of adrenergic/noradrenergic neurons. Our analysis of 894 cells resulted in 9 molecularly distinct sub-clusters (Figure 6C). By tracking the cells derived from DBH^TRAP^ and Spinal^TRAP^ samples, we identified 6 of 9 sub-clusters containing cells that were genetically tagged for *Dbh* expression (Figure 6D and table 3), whereas 6 of 9 sub-clusters contained Spinal^TRAP^ cells (Figure 6E). Both *Cartpt* and *Dbh* were present in most, but not all, sub-clusters (Figure 6F). Consistent with this result, 52 ± 4% of all *Cartpt*+ VLM neurons also expressed *Dbh*, whereas 81 ± 2% of all *Dbh*+ VLM neurons also expressed *Cartpt* (Figure 6G,H). Of note, *Slc17a6* (VGluT2) and *Slc6a2* (NET) transcripts were present in non-overlapping *Cartpt/Dbh* sub-clusters (Figure 6F). Moreover, sub-clusters expressing *Slc6a2* (n09.s06 and n09.s04) expressed *Dbh* and *Th* at relatively high levels, and *Cartpt* at relatively low levels (Figure 6F). Importantly, there were no Spinal^TRAP^ neurons in n09.s06 and n09.s04 (Figure 6E), suggesting neither noradrenergic subtypes represent A5 neurons (Souza et al., 2022a). Collectively, this indicates that there are potentially 2 subtypes of noradrenergic neurons in the VLM.

**Figure 6:**
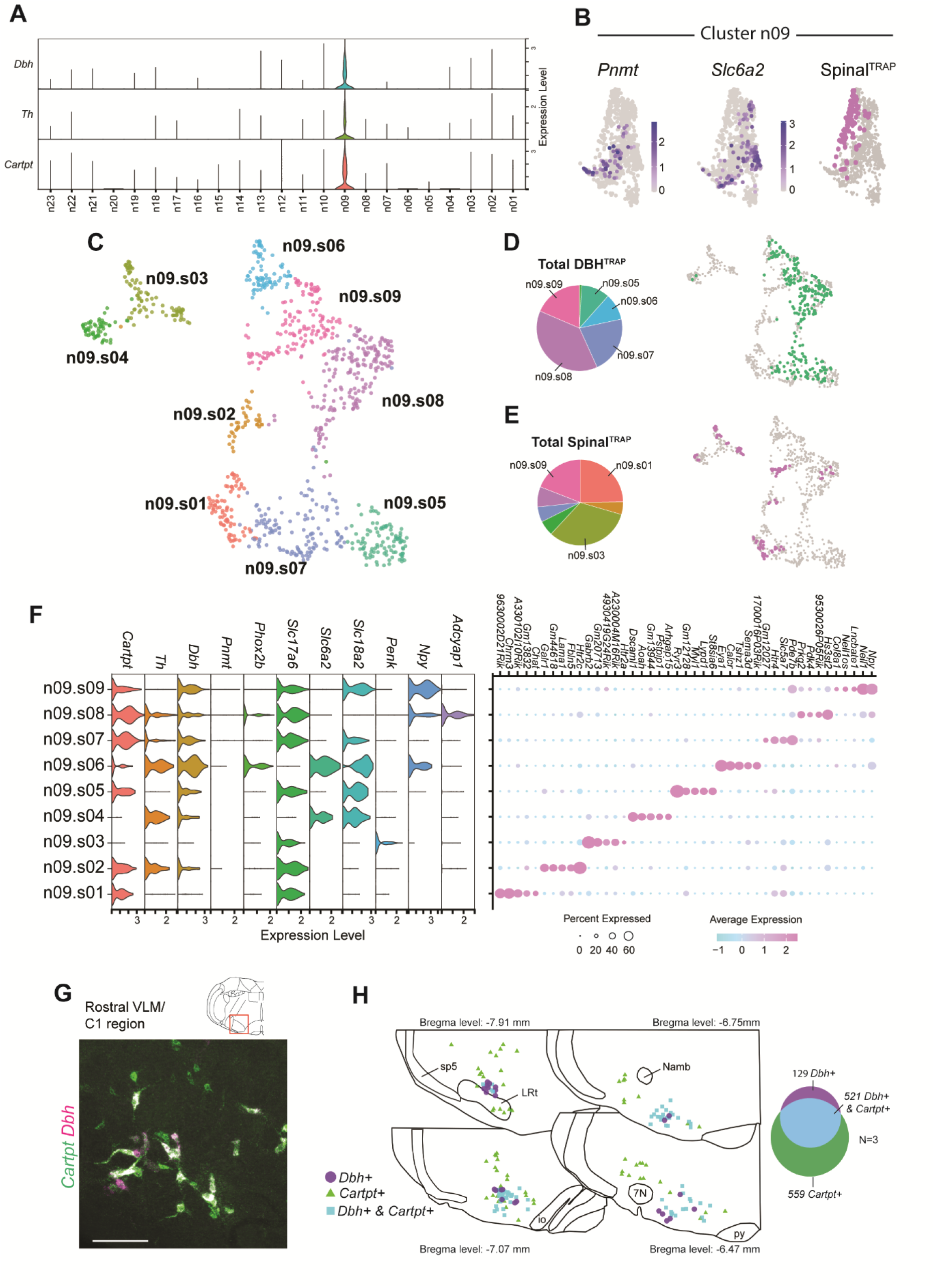
Identification of subtypes of VLM *Cartpt/Dbh* neurons. A. Violin plot of cluster marker genes for Cartpt/*Dbh* cluster n09 in ‘all-neuron’ clusters. B. Cluster n09 feature plot colored to show the distribution of *Pnmt*+, *Slc6a2*+, and Spinal^TRAP^ neurons. C. UMAP generated by sub-clustering n09 neurons based on their high-variance genes. D. UMAP from panel C is colored to show the distribution of DBH^TRAP^ neurons and a pie chart showing the proportion of DBH^TRAP^ neurons in each *Cartpt/Dbh* neuron subcluster. Values for pie chart provided in table 3. E. UMAP from panel C is colored to show the distribution of Spinal^TRAP^ samples and a pie chart showing the proportion of SpinalTRAP samples in each *Cartpt/Dbh* neuron subcluster. Values for pie chart provided in table 3 F. Expression of phenotype marker genes for C1/A1 neurons and top cluster marker genes for the *Cartpt/Dbh* neuron subclusters. G. Expression of *Dbh* and *Cartpt* in the VLM based on FISH. Scale bar-100 µm. H. Co-expression analysis of *Dbh* and *Cartpt* in the VLM based on FISH.

**Table 3.** Values for pie charts in. **figure 6**

|  | Total |  | Spinal<sup>TRAP</sup> cells |  | DBH<sup>TRAP</sup> cells |  |
| --- | --- | --- | --- | --- | --- | --- |
| Cartpt/Dbh subclusters | no. of cells | % Total | no. of cells | % Total | no. of cells | % Total |
| n09.s01 | 81 | 9.1 | 26 | 24.8 | 0 | 0.0 |
| n09.s02 | 39 | 4.4 | 5 | 4.8 | 0 | 0.0 |
| n09.s03 | 85 | 9.5 | 34 | 32.4 | 0 | 0.0 |
| n09.s04 | 59 | 6.6 | 6 | 5.7 | 2 | 0.8 |
| n09.s05 | 100 | 11.2 | 0 | 0.0 | 28 | 10.8 |
| n09.s06 | 80 | 8.9 | 0 | 0.0 | 26 | 10.0 |
| n09.s07 | 128 | 14.3 | 6 | 5.7 | 56 | 21.6 |
| n09.s08 | 186 | 20.8 | 8 | 7.6 | 99 | 38.2 |
| n09.s09 | 136 | 15.2 | 20 | 19.0 | 48 | 18.5 |
| Total | 894 |  | 105 |  | 259 |  |

Pre-sympathetic C1 neurons are known to express the gene *Slc17a6* (Stornetta et al., 2002), which encodes the vesicular glutamate transporter 2. We detected *Slc17a6* expression in the sub-clusters that expressed *Cartpt* at high levels, as well as one sub-cluster (n09.s03) that did not express *Dbh* or *Cartpt*. *Pnmt* was present in several subclusters and we observed an enrichment of *Dbh, Th* and *Slc17a6* in 5/9 subclusters (Figure 6F) consistent with descriptions of VLM adrenergic neurons (i.e. the C1 neurons). Furthermore, the co-expression of these genes (*Dbh, Th* and *Slc17a6)* across multiple *Cartpt/Dbh* sub-clusters suggests the presence of several molecularly distinct subpopulations of C1 neurons. Several neuropeptides that are known to be expressed in C1/A1 neurons in rats were enriched in *Dbh*+ subpopulations in our dataset; *Npy* was expressed in 3 clusters, one of which expressed high *Slc6a2. Npy* is enriched in C1/A1 neurons in the rat and rabbit (Blessing et al., 1987; Stornetta et al., 1999; Tseng et al., 1993). *Npy* expression was greatest in n09.s09, which contains a substantial number of Spinal^TRAP^ neurons, and expressed at lower levels in n09.s08 and n09.s06. We also found expression of the gene encoding pituitary adenylate cyclase-activating polypeptide, PACAP (*Adcyap1*), in sub-cluster n09.s08, which is expressed by spinally-projecting catecholaminergic neurons in the rostral VLM in rats (Farnham et al., 2008). Consistent with this, n09.s08 was an excitatory cluster of *Dbh*+ neurons that included Spinal^TRAP^ neurons. As such, the molecular profile of the mouse *Cartpt/Dbh* neuron subtypes identified by our dataset conforms with expectations based on anatomical studies in other species.

Importantly, our analysis also identified a number of novel genes that differentiate the sub-clusters of *Cartpt*/*Dbh* neurons (Figure 6F, right panel). To provide validation for this analysis, we examined the expression of EYA transcriptional coactivator and phosphatase 1 (*Eya1*), which was a marker gene for sub-cluster n09.06. We first assessed *Eya1* expression across all neurons in our dataset (Figure 7A), finding only sparse expression including within the *Cartpt*/*Dbh* cluster. This indicates that *Eya1* is expressed by relatively few neurons in the VLM of the adult mouse. Despite this, *Eya1* expression was highly selective for the *Cartpt*/*Dbh* sub-cluster n09.s06 (Figure 7B), which expressed high levels of *Dbh* and *Slc6a2*, and no *Slc17a6* (Figure 6F). Interestingly, the expression of *Eya1* was strongly correlated with *Slc6a2* at a single cell level, indicative of co-expression (Figure 7C). Based on this, we predicted that we would find a high degree of colocalization between the expression of *Eya1, Dbh* and *Slc6a2,* which we evaluated by FISH (n=2 mice). Indeed, *Eya1* colocalized with *Dbh* and *Slc6a2* consistently in the caudal VLM, aligning with the location of the A1 neurons, whereas *Eya1* signal was diffuse in the rostral VLM (Figure 7D-F). In the caudal VLM (i.e. the A1 region), in which virtually all *Dbh* neurons expressed detectable *Slc6a2*, we observed *Eya1* expression in 76% of *Dbh+/Slc6a2+* neurons (Figure 7G). Conversely, very few *Dbh*+ neurons in the rostral VLM (i.e. the C1 region) expressed *Slc6a2* (12/130 neurons) or *Eya1* (2/130 neurons) (Figure 7G). Interestingly, *Eya1* was also co-expressed with *Dbh* and *Slc6a2* in the A2 and A6 region, but not in the C2/C3 region, in which *Slc6a2* expression was also rare (Figure 7G,H). These observations show that *Eya1* is enriched in brainstem noradrenergic neurons, and provides a marker for a subtype of A1 cells.

**Figure 7:**
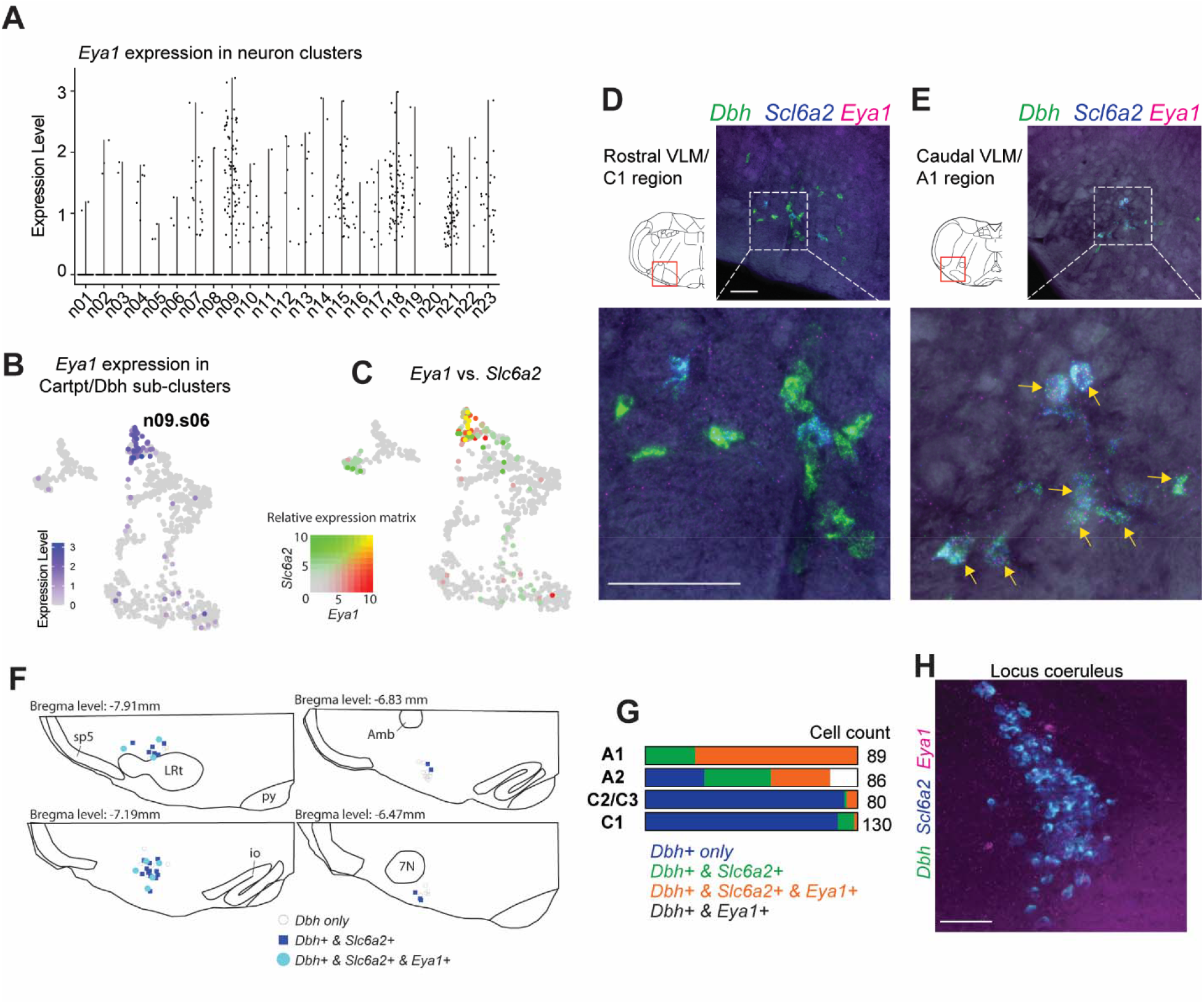
*Eya1* is enriched in VLM A1 neurons. A. Violin plot of *Eya1* expression in ‘all-neuron’ clusters. B. UMAP from Figure 6C (*Dbh*/*Cartpt* subclusters) colored to show expression of *Eya1*. C. UMAP from Figure 6C (*Dbh*/*Cartpt* subclusters) colored to show co-detection of transcripts for *Eya1* and *Slc6a2*. Co-detection of these transcripts was primarily concentrated in subcluster n09.s06. D. Expression of *Dbh, Slc6a2,* and *Eya1* in the rostral VLM (C1 region) by FISH. Scale bars-100 µm for both E. Expression of *Dbh, Slc6a2,* and *Eya1* in the caudal VLM (A1 region) by FISH. Yellow arrows in the lower panel indicate cells co-expressing *Dbh, Slc6a2,* and *Eya1.* Scale bars-100 µm for both F. Distribution and overlap of *Dbh, Slc6a2,* and *Eya1* expression in the VLM. G. Cell counts for the expression of *Dbh, Slc6a2,* and *Eya1* in medullary catecholaminergic cell groups based on 2 separate cases. H. Expression of *Dbh, Slc6a2,* and *Eya1* in the locus coeruleus (A6) by FISH. Scale bar-100 µm

To further validate our results, we examined the distribution of Hexokinase 2 (*Hk2*), a gene that encodes a kinase that phosphorylates glucose to regulate glucose metabolism, that is present in *Dbh*+ neurons based on whole mouse brain single-cell RNA-seq datasets (Langlieb et al., 2023; Yao et al., 2023). In our snRNA-seq dataset, high expression of *Hk2* was largely restricted to the *Cartpt*/*Dbh* cluster (Figure 8A) and was enriched in 4 *Cartpt*/*Dbh* neuron subclusters (n09.s01, n09.s02, n09.s07, n09.s08)(Figure 8B), of which 3 contained Spinal^TRAP^ neurons (Figure 6E and table 3). Furthermore, *Cartpt/Dbh* subclusters in which *Hk2* expression was detected exhibited low *Slc6a2* and relatively high *Slc17a6* (Figure 6F), indicative of a glutamatergic rather than noradrenergic neurotransmitter phenotype. This information suggests that *Hk2* is expressed by sub-populations of *Cartpt*/*Dbh* VLM neurons, including neurons that innervate the spinal cord. We tested this prediction by mapping *Hk2* in the VLM combined with staining for *Dbh* and retrograde tracing from the spinal cord. Based on FISH (n=2 mice), *Hk2* was expressed by 60% of *Dbh*+ neurons in the rostral VLM (132/221 neurons) (Figure 8C-E) and only 1/138 *Dbh*+ neurons in the caudal VLM (Figure 8C-E). Moreover, 59% of spinally-projecting *Dbh*+ neurons in the rostral VLM expressed detectable *Hk2* (45/76 neurons) (Figure 8C-E). Notably, we also observed *Hk2* expression in both spinally-projecting and non-spinally projecting *Dbh*+ neurons in the C2/C3 cell group regions (Figure 8C,E), whereas *Hk2* was not detectable in A1 or A2 (Figure 8E), and was not observed in A5 or A6 neurons (not shown) regardless of whether these neurons projected to the spinal cord or not. This led us to hypothesize that *Hk2* was enriched in adrenergic neurons, which are defined by the expression of *Pnmt* (Guyenet et al., 2013). Based on FISH, *Pnmt* was detected in 56% of *Hk+* neurons in the rostral VLM, whereas 78% of *Pnmt+* neurons also expressed *Hk2* (Figure 8G,E). *Hk2* also colocalized with *Pnmt* in the C2/C3 region, with 78% of *Hk2+* neurons expressing detectable *Pnmt,* and 98% of *Pnmt* neurons expressing *Hk2* (Figure 8G). *Hk2 and Pnmt* were rarely expressed by noradrenergic cells in the A1 and A2 region based on the expression of *Slc6a2* (Figure 8F,G). This collectively shows *Hk2* expression is enriched in adrenergic neurons as judged by conventional criteria irrespective of the cell’s location in the medulla.

**Figure 8:**
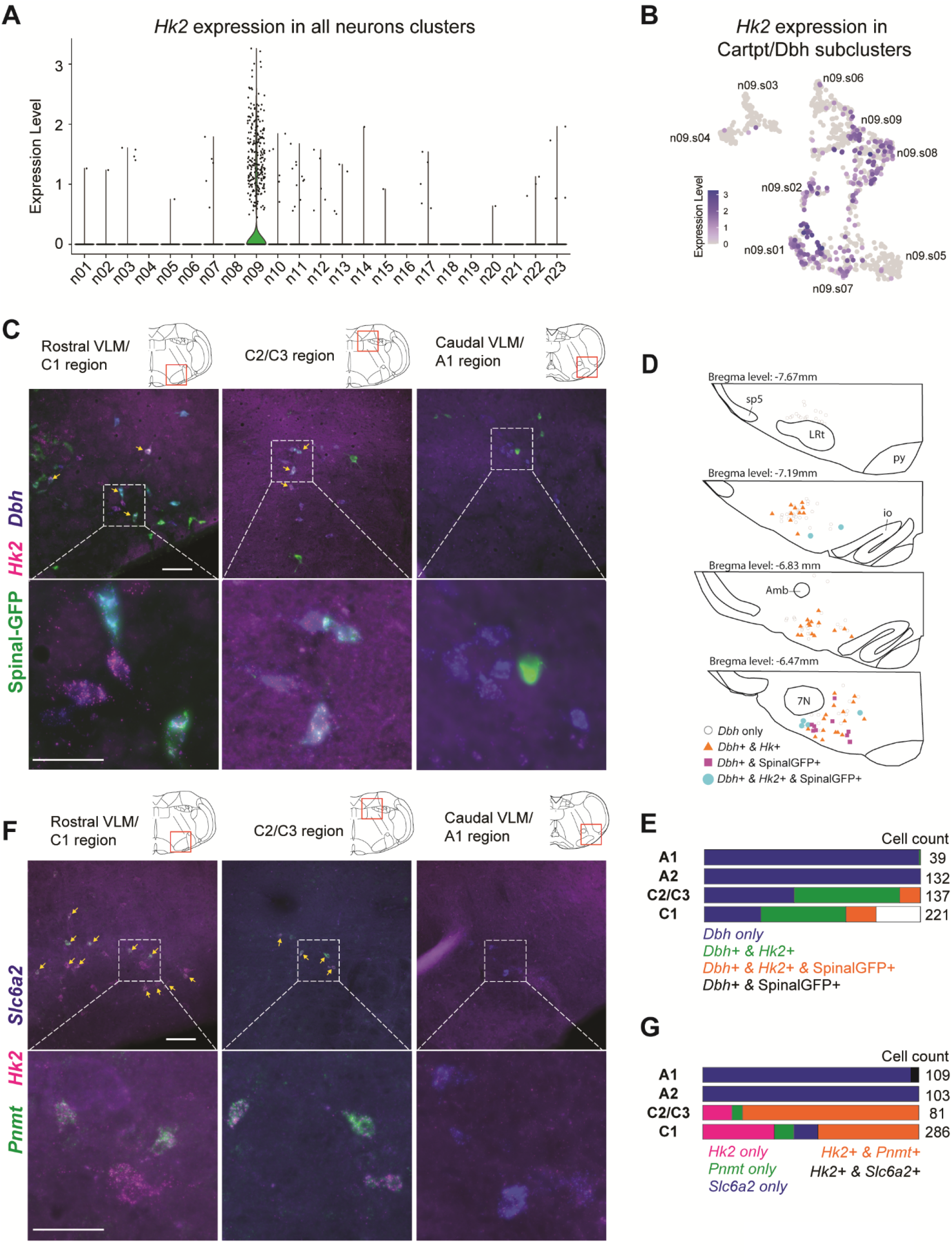
*Hk2* is enriched in VLM adrenergic neurons. A. Violin plot of *Hk2* expression in ‘all-neuron’ clusters. B. UMAP from Figure 6C (*Dbh*/*Cartpt* subclusters) colored to show *Hk2* expression. C. Expression of *Dbh* and *Hk2* by FISH, and detection of spinally-projecting neurons (Spinal-GFP) by immunofluorescence in the rostral VLM/C1 region (left panels), C2/C3 region (middle panels) and the caudal VLM/A1 region (right panels). Yellow arrows in the upper panel indicate cells co-expressing *Dbh, Hk2,* and Spinal-GFP. Scale bars-upper panels: 100 µm, lower panels: 50 µm. D. Distribution and overlap of *Dbh, Hk2,* and spinally-projecting neurons in the VLM. E. Cell counts for the expression of *Dbh, Hk2,* and spinally-projecting neurons in medullary catecholaminergic cell groups based on 2 separate cases. F. Expression of *Hk2, Pnmt,* and *Slc6a2* by FISH in the rostral VLM/C1 region (left panels), C2/C3 region (middle panels) and the caudal VLM/A1 region (right panels). Yellow arrows in the upper panel indicate cells co-expressing *Pnmt, Hk2,* and *Slc6a2*. Scale bars-upper panels: 100 µm, lower panels: 50 µm. G. Cell counts for the expression of *Dbh, Hk2,* and *Slc6a2* in medullary catecholaminergic cell groups based on 2 separate cases.

## Discussion

The study provides a detailed transcriptional atlas of VLM cell types and a dissection of the major neuronal subtypes that innervate the spinal cord. Collectively, we characterized the nuclear transcriptomes of 114,805 cells and determined specific molecular markers that can be used to identify distinct subtypes of neuronal and non-neuronal cells, infer their function, and gain genetic access to them for further investigation. Among our findings, we identify several distinct populations of spinally-projecting neurons in the VLM that are potentially involved in the control of autonomic function and motor coordination. Together our results provide new insight into molecular and cellular organization of the VLM.

### Spinal^TRAP^ Identifies the Molecular Features of Spinally-Projecting Neurons in the VLM

A goal of this study was to characterize neuronal subpopulations in the VLM that innervate the spinal cord. To achieve this we profiled the transcriptome of spinally-projecting neurons labeled with injections in the thoracic spinal cord of AAV that is efficiently internalized by axons (AAVrg-Cre) (Tervo et al., 2016). An important consideration in interpreting our results is that it was not possible to determine the effective spread of AAV injections in the spinal cord. Moreover, damage to axons *en route* to the lumbar and sacral cord may lead to transduction of neurons projecting outside of the upper thoracic segments of the spinal cord. Given these factors, it was not possible to determine the segmental targeting of Spinal^TRAP^ neurons, which will require precise projection field mapping of each cell type. Furthermore, viral tropism could affect which cells are transduced by AAVrg, which could skew the proportions of each spinally-projecting population based on our dataset. Nevertheless, our analysis identifies novel spinally-projecting clusters in the VLM as well as provides the first detailed transcriptome for bulbospinal *Cartpt/Dbh* and 5HT neuron types, which were expected be present in the Spinal^TRAP^ populations (Bowker et al., 1981; Tucker et al., 1987). This was despite reports that AAVrg is inefficient at transducing serotonergic and noradrenergic cell types (Ganley et al., 2021; Tervo et al., 2016). This discrepancy may be explained by the higher efficiency of the Cre-reporter expression systems compared to promoter-based expression systems when expressing H2B-mCherry or a similar reporter. The novel clusters of spinally-projecting neurons in our dataset included 1 excitatory cluster and 2 inhibitory clusters. Two of the bulbospinal populations (n06 and n20) may represent partially overlapping inhibitory and excitatory reticulospinal cells that are involved in locomotion and limb control (Leiras et al., 2022; Ruder and Arber, 2019). Hence, we expect the novel genes identified in this study will be useful to target brainstem locomotor circuits.

### Molecular Characterization of Adrenergic/Noradrenergic Neurons in the VLM

Monoaminergic neurons in brainstem were characterized in the 1960s and ‘70s by Dalhstrom, Fuxe, and others (Dahlstrom and Fuxe, 1964; Fuxe et al., 1970; Hökfelt et al., 1974). Since then, the expression of monoaminergic cell markers for noradrenergic, adrenergic and serotonergic neurons in the VLM has provided an essential neuroanatomical handle for exploring the integrative function of this brain region. Catecholaminergic neurons in the VLM are a highly conserved group of cells classified as C1 or A1 based primarily on the expression of *Pnmt* and their relative location in the medulla (Blessing et al., 1978; Dormer et al., 1993; Halliday and McLachlan, 1991; Howe et al., 1980; Minson et al., 1990; Reiner and Vincent, 1986, 1986; Ruggiero et al., 1992). In addition, experimental studies indicate that subpopulations of C1 and A1 cells exist based on efferent connectivity (Stornetta et al., 1999; Tucker et al., 1987) and gene expression (Guyenet et al., 2013; Halliday and McLachlan, 1991; Phillips et al., 2001; Pilowsky and Goodchild, 2002). Our data indicate there are 7 molecularly distinct subpopulations of *Dbh* neurons, including 5 adrenergic subtypes and 2 noradrenergic subtypes. We histologically verified two novel markers of the adrenergic and noradrenergic neurons, one generated by our own data (*Eya1*) and the other (*Hk2*) identified in other studies (Langlieb et al., 2023; Yao et al., 2023). This study indicates that *Eya1* is a unique marker for a subpopulation of noradrenergic neurons in the caudal VLM and also other noradrenergic subpopulations in the dorsal medulla and pons, but not adrenergic neurons. In contrast, *Hk2* was expressed in 4 excitatory *Cartpt/Dbh* sub-clusters and neither of the noradrenergic sub-clusters, suggesting this gene was expressed primarily by glutamatergic/adrenergic cells. Consistent with this, *Hk2* expression was found primarily in *Dbh+* neurons in the rostral VLM, including both bulbospinal and non-bulbospinal neurons, as well as as majority of *Pnmt+* neurons in the rostral VLM and C2/C3 regions. Hence, *Hk2* expression appears to be a novel and fairly selective positive selection criterion for adrenergic neurons in the medulla. In addition to validating our snRNA-seq dataset, our identification of *Eya1* and *Hk2* reaffirms the notion that molecularly distinct adrenergic and noradrenergic neurons are spatially intermixed in the VLM, emphasizing the need for precise molecular and genetic approaches to dissect the function of each subtype.

### Spinal^TRAP^ Identifies Putative Marker Genes for Non-Catecholaminergic Pre-Sympathetic Neurons in the Rostral VLM

In rats, adrenergic neurons represent a majority of neurons in the rostral VLM that project to sympathetic preganglionic neurons in the spinal cord, accounting for up to 70% of the total pre-sympathetic neurons in this region of the brain (Lipski et al., 1995; Schreihofer and Guyenet, 1997). These cells have a well-established role in the sympathetic control of blood pressure (Souza et al., 2022b). The remainder of the pre-sympathetic population in the VLM in rats are non-catecholaminergic excitatory neurons, although a distinguishing marker for the ‘non-C1 neurons’ remains elusive. In our dataset, neurons expressing adrenergic/noradrenergic genes (*Dbh*, *Th*) also expressed the neuropeptide gene *Cartpt,* similar to what has been found in rats (Burman et al., 2004; Dun et al., 2002; Koylu et al., 1998). Moreover, Burman et al., (2004) suggested that *Cartpt* identifies pre-sympathetic VLM neurons involved in cardiovascular function. Interestingly, by sub-clustering the *Cartpt/Dbh* cells we identified a sub-type of glutamatergic neurons that express *Cartpt* and are largely devoid of catecholamine cell markers (neuron subcluster n09.s01). Many cells in this subcluster were Spinal^TRAP^ cells, and therefore, this cluster could represent non-catecholaminergic pre-sympathetic neurons in the VLM. Another candidate for non-catecholaminergic pre-sympathetic neurons is subcluster n09.s03 as it contained a substantial number of Spinal^TRAP^ cells and expressed *Penk* (Stornetta et al., 2001), but no detectable *Cartpt* or *Dbh*. Future studies involving genetically-targeted functional experiments using novel markers for sub-cluster n09.s01 and n09.s03 could offer new insight into the identity of VLM neurons that underlie the sympathetic regulation of blood pressure.

### Molecular Characterization of Serotonergic Neurons in the Parapyramidal Region

Serotonergic neurons in the parapyramidal region are implicated in thermoregulation through sympathetic control of cutaneous blood flow, brown adipose tissue thermogenesis, cardiac and sudomotor function (Bamshad et al., 1999; Cano et al., 2003; Dampney and McAllen, 1988; McAllen, 1986; Nakamura et al., 2004; Pelaez et al., 2002), as well as respiratory functions (Brust et al., 2014; da Silva et al., 2010; Ribas-Salgueiro et al., 2005, 2005). This study provides a detailed transcriptomic analysis of 5HT neurons in the parapyramidal region, a region that has been overlooked by previous studies. We discovered marker genes (*Cpa6*, *Cpne7*) that define two broad classes of parapyramidal 5HT neurons, as well as several genes that distinguished 6 putative sub-types within these classes. Furthermore, genes that are expressed in distinct types of 5HT cells in other regions of the brain were also evident in different subtypes of 5HT neurons in the the parapyramidal region (for example, *Tacr1*, *Gad1/2*, *Calcr*, *Adra1a*, *Trh*) (Huang et al., 2019; Okaty et al., 2015; Schneeberger et al., 2022; Xiao et al., 2021). This suggests that parapyramidal 5HT neurons are composed of molecularly distinct subpopulations, similar to 5HT groups in other brain regions.

### Identification of putative markers for excitatory and inhibitory respiratory-related neurons

The VLM contains the neural network that generates breathing (Del Negro et al., 2018; Feldman et al., 2003). The classification of VLM neurons that generate the rhythm and pattern of breathing is principally based on their firing properties in relation to the respiratory cycle (Richter and Smith, 2014; Smith, 2022). However, some studies have identified genes that are expressed by respiratory-related neurons with varying degrees of selectivity (Bachmutsky et al., 2020; Cui et al., 2016; Gray et al., 1999; Stornetta, 2008; Stornetta et al., 2003b). We leveraged this information to identify putative respiratory-related clusters in our dataset (n05 and n12). These clusters appear to represent two classes of neurons distinguished from each other by their neurotransmitter phenotype (i.e., excitatory and inhibitory), and from other cells in the dataset by the differential expression of specific genes, such as *Glra3* and *Gpc3*. Future studies are needed to confirm that the genes identified in cluster n05 and n12 label respiratory-related neurons. This study also shows that *Trpc4,* a non-selective calcium-permeable cation channel regulated by G_q_ and G_i/o_ G-protein-coupled receptor signaling (Schaefer et al., 2000; Thakur et al., 2016; Tian et al., 2022), is expressed at relatively high levels in both n05 and n12 compared to other neurons in our dataset. *Trpc4* expression has been reported in functionally identified auto-rhythmic respiratory neurons in the Pre-Bötzinger Complex of neonatal mice (Kallurkar et al., 2022), providing additional support for our conclusion that n05 and n12 contains respiratory-related neurons. Given the abundance of *Trpc4* expression in n05 and n12, we propose that the activity of TRPC4 in VLM respiratory neurons integrates the co-activation of G_q_ and G_i/o_ signaling, and as such, contributes to the state-dependent control of respiratory motor output (Ramirez et al., 2012).

### Conclusions and Implications for Future Studies

Our study highlights the diversity of cell types in the VLM, many of which overlap in anatomical distribution, and provides a valuable resource for future studies involving the VLM. Leveraging this data, which can be explored using the Broad Institute Single Cell Portal, will facilitate a rigorous delineation of cells involved in the regulation of autonomic, respiratory, and motor function. This dataset also provides a snapshot of the transcriptional state of the VLM cells in healthy adult mice that can be referenced to identify and characterize signaling pathways that may be affected in diseases that involve changes in VLM function, such as neurodegenerative disease (Benarroch, 2019) and neurogenic hypertension (Guyenet et al., 2020), or through comparisons with genome-wide association studies. Overall these data provide a basis for generating novel hypotheses concerning VLM function and offer the means to test them.

## Methods

### Animals

All experiments were conducted in accordance with the National Institutes of Health’s *Guide for the Care and Use of Laboratory Animals* and approved by the University of Virginia Animal Care and Use Committee (protocol #4312). Animals were group-housed wherever possible and maintained under a 12:12 light: dark cycle at 23°C-24°C with water and food provided *ad libitum*.

We obtained H2B-TRAP mice (Roh et al., 2017) (RRID:IMSR_JAX:029899) as a gift from Evan Rosen and Linus Tsai (Beth Israel Deaconess Medical Center, Harvard Medical School). H2B-TRAP mice express mCherry-tagged nuclear H2B protein (H2B-mCherry) and an eGFP-tagged ribosomal L10 protein from the Rosa26 genomic locus upon removal of a translational stop cassette by Cre recombination. Dbh-Cre mice (strain STOCK Tg(Dbh-cre)KH212Gsat/Mmcd) (RRID:MMRRC_032081-UCD) procured from the Mutant Mouse Regional Resource Centers were maintained as Cre hemizygous (Dbh^Cre/+^). Dbh^Cre/+^ mice were crossed with H2B-TRAP homozygote mice to generate DBH^TRAP^ mice. Dbh^+/+^ Cre-negative mice hemizygous for H2B-TRAP were injected with AAVrg-Cre in the spinal cord to generate Spinal^TRAP^ mice. For anatomical tract-tracing studies, we used R26-loxSTOPlox-L10-GFP Cre-reporter mice acquired from The Jackson Laboratory (strain #024750, RRID:IMSR_JAX:024750) and bred as heterozygotes with C57BL/6J mice in house.

### Single-Nuclei RNA-Seq Library Generation

To avoid stress- and anesthesia-related changes in gene expression, we rapidly decapitated unanesthetized mice upon scruffing and extracted their brains. Each sample batch consisted of tissue from 5-8 mice of each sex. Each brain was placed in ice-cold Earl’s Balanced Salt Solution (catalog # E2888; Sigma-Aldrich; St. Louis, MO, U.S.A.) or Hibernate-A (catalog # A1247501; Thermo Fisher Scientific; Waltham, MA, U.S.A.) for 2-5 min, transferred dorsal surface up to a chilled brain matrix (catalog # SA-2175; Roboz; Gaithersburg, MD, U.S.A.), and sectioned in the coronal plane at 1 mm intervals. Keeping the tissue chilled throughout, we microdissected RVLM tissue with a scalpel blade on an ice-filled petri dish. We then processed brain tissue into cell nuclei suspensions using one of two protocols, as described below.

For sample batches 1 and 2, we transferred the dissected tissue to an RNase-free microcentrifuge tube, froze it in a slurry of dry ice and methanol, and then stored it in a -80□ for less than a month. After thawing a batch of tissue samples on ice, we isolated cell nuclei from the thawed tissue using a previously described hypotonic lysis protocol (Matson et al., 2018), modified as follows: trituration with 5 Pasteur pipettes, starting with an unpolished pipette, followed by a series of four pipettes with tips fire polished to progressively smaller diameters; 5 to 10 strokes for each size pipette, or until the tissue passed smoothly through the pipette; and filtered resuspended nuclei with 20 um strainer (pre-separation filter; catalog # 130-101-812; Miltenyi Biotec; Bergisch Gladbach, Germany). We fluorescently labeled nuclei with propidium iodide and then quantified them with a DeNovix CellDrop automated cell counter. We then proceeded with the Chromium Next GEM Single Cell 3□ v3.1 kit (catalog # PN-1000128 or PN-1000121; 10X Genomics; Pleasanton, CA; U.S.A.) according to the manufacturer’s instructions (CG000204 Rev D; 10X Genomics), tagging each sequencing library with a unique 10X Sample Specific Index (catalog # PN-1000213; 10X Genomics) for sequencing libraries in multiplex.

For sample batches 3 through 6, which involved fluorescence-activated cell sorting (FACS), we isolated cell nuclei from preserved tissue samples based on a published protocol (Habib et al., 2016; Todd et al., 2020). Briefly, we transferred the microdissected tissue to RNase-free microcentrifuge tubes containing ice-cold RNAprotect reagent (catalog # 76104; Qiagen; Venlo, The Netherlands). We stored the tissue samples in RNAprotect at 4□ overnight, then removed the liquid and transferred the preserved tissue samples to -80 □ storage for no more than 3 months. When ready to proceed, we thawed a batch of tissue samples on ice, dounce homogenized them in 1 mL of homogenization buffer 25 times with pestle A and 25 times with pestle B (Kimble Kontes Dounce Tissue Grinder, DWK Life Sciences, catalog #885300-0001), then separated the nuclei from debris by density gradient centrifugation according to the protocol (Habib et al., 2016). We filtered the resuspended nuclei through a 20 um cell strainer (Miltenyi Biotec pre-separation filter), fluorescently labeled DNA with NucBlue Live ReadyProbes Reagent (catalog # R37605, Thermo Fisher Scientific), and sorted the nuclei on a Becton Dickenson Influx cell sorter equipped with an 85 µm nozzle and set to purity mode. All sorts were performed with Specialist in Cytometry (SCYM; ASCP) certified technical assistance at the University of Virginia Flow Cytometry Core. To separate nuclei from non-nucleated debris, we gated for events with high relative intensity of NucBlue fluorescence. We then gated the nuclei by forward scatter area vs. side scatter area to eliminate large aggregates, followed by forward scatter area vs. forward scatter height to select for singlets. Finally, from the single nuclei, we selected H2B-mCherry+ and -negative nuclei based on their high and low relative fluorescence, respectively, with 561 nm excitation and 610/20 nm collection filter. We separated H2B-mCherry+ and -negative nuclei singlets into low-bind microcentrifuge tubes, each containing 18.8 uL of RT Reagent B (from Chromium Next GEM Single Cell 3□ v3.1 kit). We added the remainder of the 10X master mix reagents to each tube of sorted nuclei, before proceeding with the kit according to the manufacturer’s instructions (CG000204 Rev D; 10X Genomics), tagging each sequencing library with a unique 10X Sample Specific Index (catalog # PN-1000213; 10X Genomics) for sequencing libraries in multiplex.

### Single-Nuclei RNA-Seq Data Generation, Processing, and Analysis

We sequenced the single-nuclei RNA-seq libraries on an Illumina Next-Seq 550 with high output, 75-cycle kits (catalog # 20024906; Illumina; San Diego, CA, U.S.A.). After sequencing each single-nucleus library to a targeted minimum of 20,000 reads per nucleus, we demultiplexed the sequencing reads, aligned them to the mouse genome mm10 with CellRanger v5.0 (10X Genomics) including intronic reads with the command --use-introns, and quantified gene-level (feature) expression values for each cell-specific barcode as a feature-barcode matrix. We then processed each feature-barcode matrix with CellBender (v0.2.2; parameters provided in Supplementary Figure 1) to remove ambient RNA reads and random barcode swapping from the dataset, with the exception of batches 1 and 2 where the lack of a clear inflection point in the barcode rank plot undermined CellBender’s performance.

We analyzed the data with Seurat v.4.3.0.1 software package in R (version 4.0.2) using default Seurat parameters unless otherwise specified. Notably, the H2B-mCherry+ cell nuclei from A5 noradrenergic neurons were excluded from the analysis because of low yield, preventing a robust analysis. Across all samples, we excluded cells whose transcriptomes contained greater than 5% mitochondrial transcripts. To remove putative doublet samples, the upper quartile (Q3) and the interquartile range (IQR) of the number of genes and the number of UMIs in samples in each batch were calculated. Within each batch, the formula Q3 + 3*IQR was used to calculate the upper cutoff values for the number of genes and the number of UMIs, and samples exceeding these values were excluded from further analysis. The lower cutoff for the number of genes was different per batch and was chosen based on each batch’s distribution of this quality metric. For each batch we log-normalized the data; selected the 2,000 most variable genes (feature selection); scaled expression of each gene; and used Principal Component Analysis (PCA) for linear dimensionality reduction of the transcriptomes in high-variance gene space. Reciprocal Principal Component Analysis (RPCA) based integration was then used to correct for technical differences between batches. This integration created a new integrated assay, for which we then scaled the expression of each gene; used PCA for linear dimensionality reduction of the transcriptomes in a high variable gene space; clustered the cells using the Louvain algorithm, based on the Euclidean distance in the PCA space comprising the first 30 principal components (PCs) and with a resolution value of 0.5; and performed non-linear dimensionality reduction by Uniform Manifold Approximation and Projection (UMAP) for visualization of the dataset in two dimensions.

To match each sample to a known cell type, we checked each cluster’s expression for cell type marker genes (Table 4) and annotated them accordingly. We identified marker genes for each cluster using the Seurat FindAllMarkers function, returning only positive markers but with otherwise default parameters, which used a Wilcoxon rank-sum test and Bonferonni-corrected significance threshold to compare gene expression of cells within a given cluster against all other cells.

**Table 4.** Cell type identity genes.

| Known Cell Types | Marker Genes |
| --- | --- |
| Oligodendrocytes | *Olig1, Olig2, Olig3* |
| Polydendrocytes | *Olig1, Cspg4* |
| Astrocytes | *Gfap, Slc13a3, Slc1a2, Agt* |
| Microglia | *Tmem119, Cx3cr1* |
| Endothelial Cells | *Slco1c1* |
| Neurons | *Rbfox3, Syp, Syn1* |
| Olivary Neurons | *Rbfox3, Syp, Syn1, Foxp2* |
| Vascular leptomeningeal cells | *Ranbp3l* |
| Pericytes | *Kcnj8, Abcc9* |
| Choroid Plexus Cells | *Kl, Folr1* |

We identified clusters representing cell doublets (*i.e*., transcriptome from more than one cell) based on their (1) co-expression of cell-type specific marker genes and (2) otherwise lacking specific marker genes, and excluded these doublet clusters from further analysis. We identified ambiguous clusters, likely representing cells with degraded or contaminated transcriptomes, based on their lack of specific marker genes. After removing doublet and ambiguous clusters, we repeated the post-integration data processing with the same parameters. For each subsequent cell-type level analysis (astrocytes, all neurons, 5HT neurons, and *Cartpt/Dbh* neurons), we repeated this pipeline of separately processing each batch, RPCA integration, followed by re-processing the new integrated assay and identification of marker genes with the Seurat FindAllMarkers function. In these analyses, if doublet or ambiguous clusters were removed after the first post-integration analysis, a second round of this pipeline was performed until all doublet and ambiguous clusters were removed. This iterative workflow and parameters are illustrated in Supplementary Figure 1.

To further analyze the neurons in our dataset, we subsetted the clusters labeled as neurons based on their expression of classical neuron marker genes. To ensure only high-quality neuron samples were included in this subsequent analysis, cells within these clusters that did not individually express *Syn1, Rbfox3,* or *Syp* were excluded. Neurons in which we detected expression of the oligodendrocyte marker gene *Prr5l* were suspected as contaminated or doublets and were also excluded.

### Perfusion and Histology

Mice were deeply anesthetized with a mixture of ketamine (100 mg/kg) and dexmedetomidine (0.2 mg/kg) given i.p. and perfused transcardially with 4% paraformaldehyde, pH 7.4 in 100 mM phosphate buffer or pH-buffered 10% formalin, pH 7.0. Brains were removed and post-fixed in the same fixative for 12-24 hrs. at 4°C. Brains were sectioned (30-50 µm) on a vibratome (VT-1000S, Leica Biosystems, Deer Park, IL, USA), and sections were stored in cryoprotectant (30% ethylene glycol (v/v), 20% glycerol (v/v), 50% 100mM phosphate buffer, pH 7.4) at −20°C. Immunohistochemistry was performed on free-floating sections at room temperature unless noted otherwise. Serial 1-in-3 sections were rinsed, then blocked in a solution containing 100 mM Tris, 150 mM saline, 10% horse serum (v/v), 0.1% Triton-X (v/v) and incubated with primary antibodies for 60 min at room temperature then 4°C overnight. Sections were subsequently rinsed and then incubated with secondary antibodies for 60 min at room temperature and rinsed again before mounting on slides. Slides were mounted with ProLong Gold antifade reagent with DAPI (P36931, Thermo Fisher Scientific).

Multiplex fluorescent *in situ* hybridization was performed using RNAscope (V1 kit, Advanced Cell Diagnostics, Newark, CA, USA). Serial 1-in-3 sections were washed in RNase(ribonuclease)-free phosphate buffered saline, mounted on charged slides, dried overnight, and processed according to the manufacturer’s instructions. When necessary, immunohistochemistry was performed on the slide after the RNAscope procedure using antibodies against mCherry or GFP along with their respective secondary antibodies. Immediately following the RNAscope procedure, sections were rinsed and then incubated in blocking solution for 10 min followed by incubation in primary at room temperature for 60 min, rinsed and incubated in secondary antibodies for 30 min, rinsed and then dried overnight before cover slipping. Slides were cover slipped with Prolong Gold antifade mountant with DAPI (P3693, Thermo Fisher Scientific). Primary and secondary antibodies and FISH probes used are detailed in Table 5 and 6 respectively.

**Table 5.** Antibodies used for immunohistochemistry.

| Primary Antibodies |  |  |  |  |  |
| --- | --- | --- | --- | --- | --- |
| Host | Target | Dilution | Vendor | Catalog | RRID |
| Chicken | GFP | 1:1k | Aves Labs | GFP-1020 | AB_10000240 |
| Sheep | TH | 1:1k | MillipreSigma | AB1542 | AB_90755 |
| Rabbit | DsRed | 1:1k | Takara Bio | 632496 | AB_10013483 |
| Secondary Antibodies |  |  |  |  |  |
| Host | Target | Dilution | Fluorophore | Catalog | RRID |
| Donkey | Chicken IgY | 1:500 | 488 | 703-545-155 | AB_2340375 |
| Donkey | Rabbit IgG | 1:500 | Cy3 | 711-165-152 | AB_2307443 |
| Donkey | Sheep IgG | 1:500 | 647 | 713-606-147 | AB_2340752 |
| Secondaries purchased from Jackson ImmunoResearch. |  |  |  |  |  |

**Table 6.**
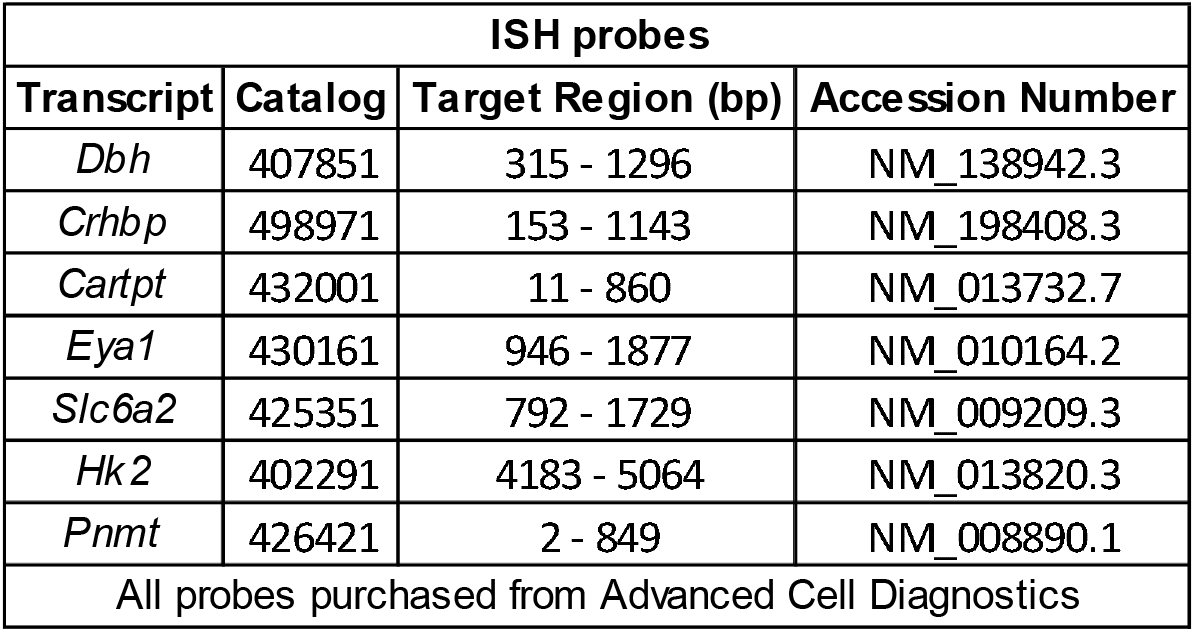
Probes used for fluorescent in-situ hybridization.

| ISH probes |  |  |  |
| --- | --- | --- | --- |
| Transcript | Catalog | Target Region (bp) | Accession Number |
| *Dbh* | 407851 | 315 - 1296 | NM_138942.3 |
| *Crhbp* | 498971 | 153 - 1143 | NM_198408.3 |
| *Cartpt* | 432001 | 11 - 860 | NM_013732.7 |
| *Eya1* | 430161 | 946 - 1877 | NM_010164.2 |
| *Slc6a2* | 425351 | 792 - 1729 | NM_009209.3 |
| *Hk2* | 402291 | 4183 - 5064 | NM_013820.3 |
| *Pnmt* | 426421 | 2 - 849 | NM_008890.1 |
| All probes purchased from Advanced Cell Diagnostics |  |  |  |

### Microscopy, Imaging and Cell Mapping

Neuronal mapping was conducted using Neurolucida software (MBF Bioscience, Williston, VT, USA) with an AxioImager M2 microscope (Carl Zeiss Microscopy, White Plains, NY, USA). Digital images were acquired in grayscale using a Hamamatsu C11440 Orca-Flash 4.0LT digital camera (Hamamatsu). Max projections of z-stack images were exported (8-bit) and further image modifications were performed in Fiji software (Schindelin et al., 2012). Representative images were pseudo-colored and optimized for presentation, brightness and contrast was adjusted equally in all pixels of the image. Cell counting was performed manually and unblinded using a 1-in-3 series that was subsequently aligned to Paxinos and Franklin’s 4th edition (Paxinos and Franklin, 2012). In Figure 8 & 9, cell counts are reported for the rostral, caudal VLM, A2, and C2/C3 region. Rostral VLM was considered to be a region within 500 µm caudal of the facial nucleus (bregma level: -6.47 to -6.83 mm). Caudal VLM was considered to be a region within 500 µm rostral of the spinal decussation (bregma level: -7.55 to -8.05 mm). A2 neurons were counted in the ventrolateral nucleus of the solitary tract (bregma level: -7.55 to -7.67 mm); counts were based on neurons expressing intense *Dbh* or *Slc6a2*, considered to be A2 neurons. The C2/C3 region was considered the region of the dorsal medial medulla between (bregma level: -6.75 to -6.95 mm). To generate these counts of cells in the aforementioned regions, all cells expressing the gene/genes of interest were counted in 2 or 3 adjacent sections in a 1 in 3 series. Cells were considered to express a gene of interest if the cell contained at least 5 particles of fluorescent signal that were not in multiple fluorescent channels.

## Acknowledgements

This work was supported by National Institute of Health (NIH) R01 HL148004 to SBGA; and R01 HL153916, Pathway to Stop Diabetes Initiator Award 1-18-INI-14, and UVA 3Cavaliers grant to JNC.

## Author contributions

Conceptualization and Methodology: DCS., RJAF, JNC, SBGA

Investigation: DCS, RJAF, MEC, JNC, SBGA

Data curation: DCS, RJAF., GMPRS, MJ, JNC, SBGA

Writing – original draft preparation: DCS, DSS, JNC, SBGA

Writing – review & editing: all authors.

Visualization: DCS, DSS, JNC, SBGA

Supervision: JNC, SBGA

Funding acquisition: JNC, SBGA

## Conflict of interest

The authors declare no competing interests.

---

**Supplemental Figure 1:**
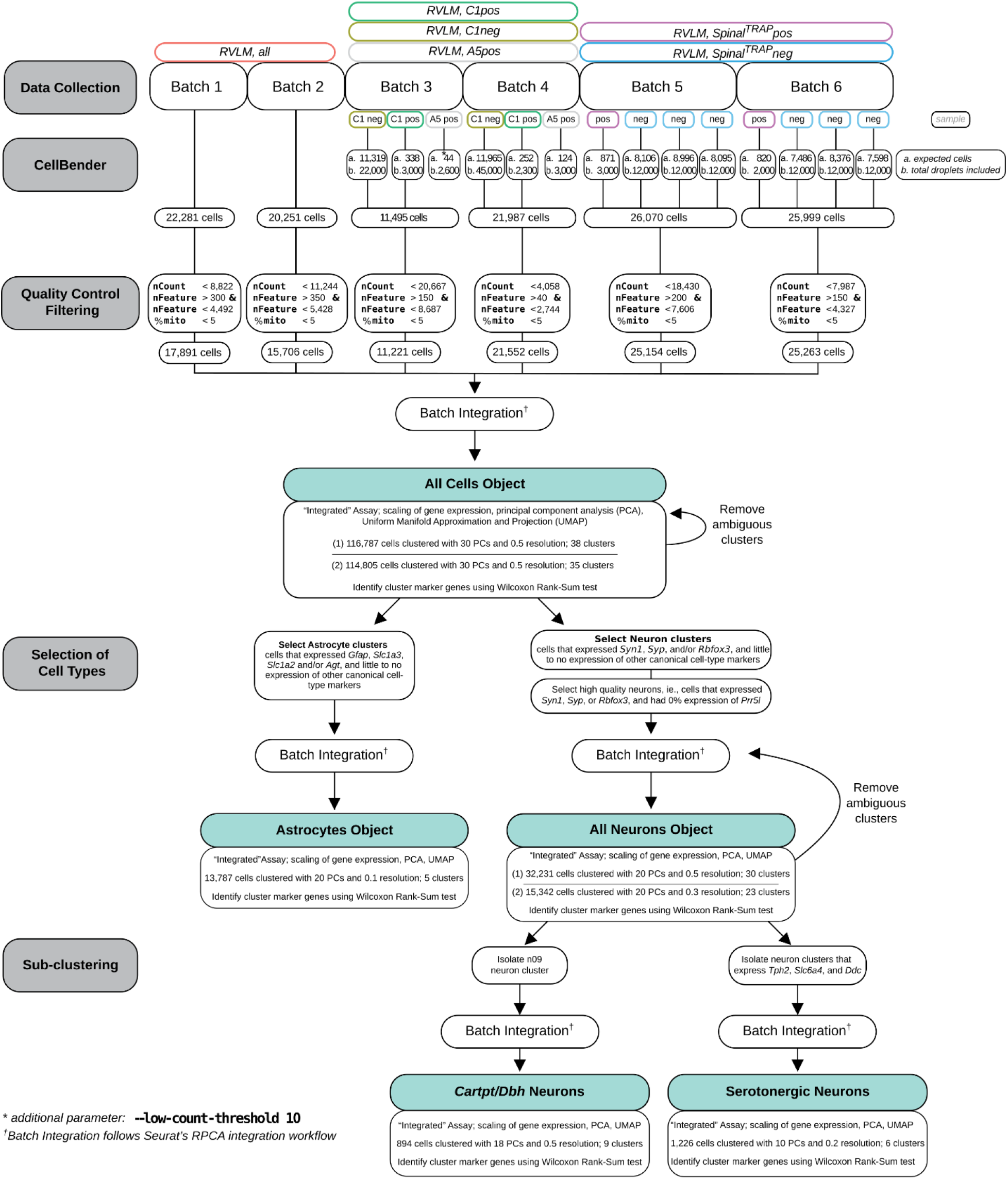
Data processing workflow and parameters.

**Supplemental Figure 2:**
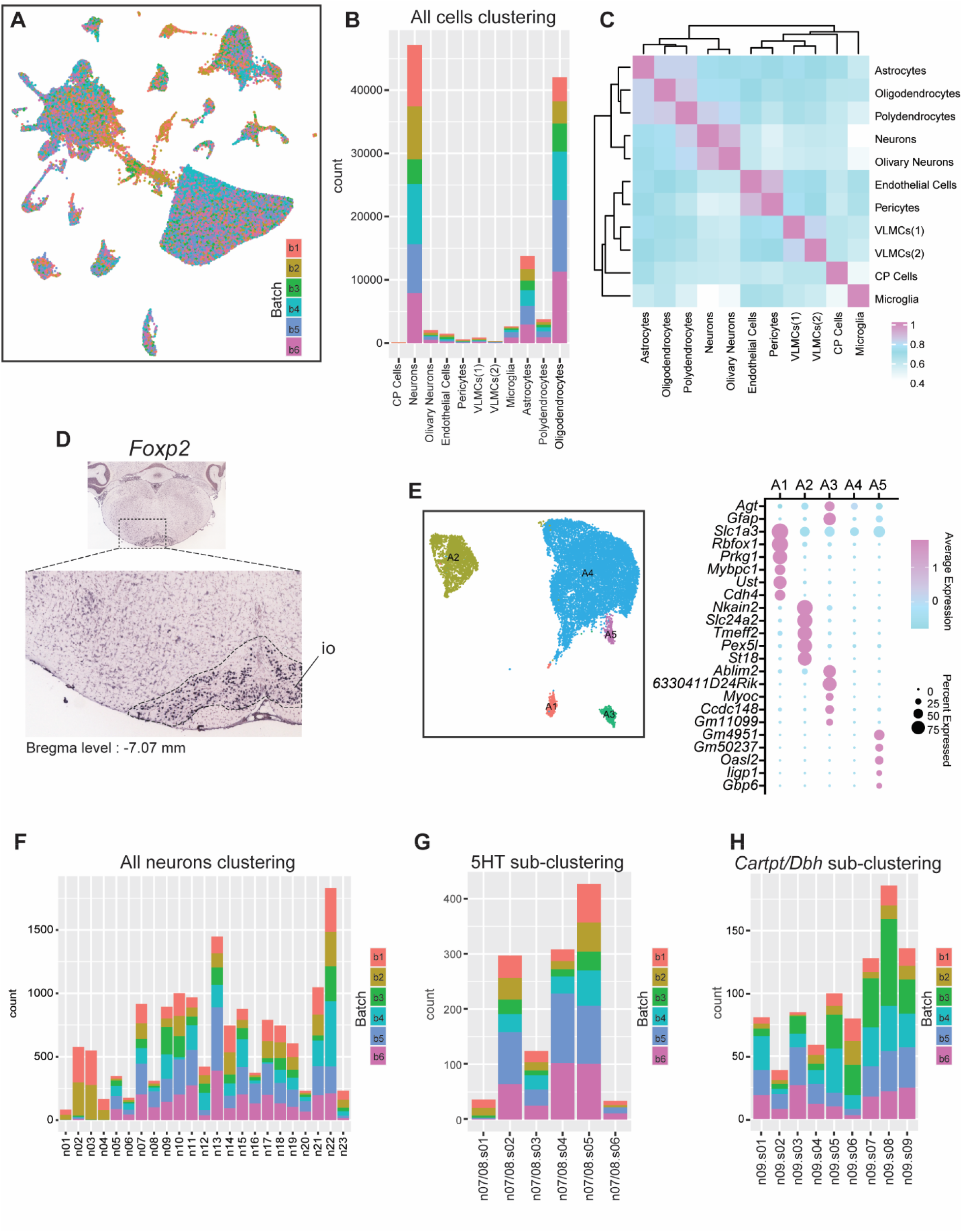
Data integration and subtypes of astrocytes. A. UMAP of all cells colored by sample batch. B. Batch composition per-cluster for all cells. C. Cluster correlation matrix for all cells. D. Expression of *Foxp2* from Allen Mouse Brain Atlas (https://mouse.brain-map.org/experiment/show/72079884). Note the intense expression in the inferior olive. E. UMAP of astrocyte subtypes and dot plot of differentially expressed genes in astrocyte subtypes F. Batch composition for each cluster for neurons. G. Batch composition for each sub-cluster for 5HT neurons. H. Batch composition for each sub-cluster for *Cartpt/Dbh* neurons.

## Notes

https://abbott-lab.com/single-nuclei-rnaseq-datasets/

### Competing Interest Statement

The authors have declared no competing interest.

